# Next-Generation Multiplexed Targeted Proteomics Quantifies Post-Translational Modifications, Compound-Protein Interactions, and Disease Biomarkers with High Throughput

**DOI:** 10.1101/2025.09.10.675380

**Authors:** Steven R. Shuken, Geordon A. Frere, Charlotte R. Beard, Jesse D. Canterbury, Nathan R. Zuniga, Brandon M. Gassaway, Shane L. Dawson, Kean Hean Ooi, João A. Paulo, M. Windy McNerney, Steven P. Gygi, Qing Yu

## Abstract

The GoDig platform enables sensitive, multiplexed targeted pathway proteomics without manual scheduling or synthetic standards. Here we present GoDig 2.0, which increases sample multiplexing to 35-fold, improves time efficiency and reduces scan delays for higher success rates, and allows flexible spectral and elution library generation from different mass spectrometry data types. GoDig 2.0 measures 2.4× more targets than GoDig 1.0, quantifying >99% of 800 peptides in a single run. We compiled a library of 23,989 human phosphorylation sites from a phosphoproteomic dataset and used it to profile kinase signaling differences across cell lines. In human brain tissue, we established a hyperphosphorylated tau assay including pTau127, revealing potential biomarkers for Alzheimer’s disease. We also quantified diglycyl-lysine peptides to assess polyubiquitin branching. Finally, we built a library of 20,946 reactive cysteines and profiled covalent compound-protein interactions spanning diverse pathways. GoDig 2.0 enables high-throughput analyses of site-specific protein modifications across many biological contexts.

## INTRODUCTION

The detection and measurement of proteins and their modifications is a powerful approach in biological research. Whereas the untargeted detection and quantification of thousands of proteins has become routine, these approaches are often lacking for certain proteins of interest due to limitations of scope, sensitivity, dynamic range, run-to-run overlap/completeness, and/or quantitative accuracy and precision.^1^ Targeted approaches can address these issues,^2^ and targeted mass spectrometry (MS) has advantages over antibody-based techniques: hundreds of proteins can be feasibly quantified in a single run with high quantitative accuracy and precision, and peptides’ ambiguities are well defined, allowing the development of highly specific “proteotypic” peptide-based mass spectrometric assays.^3,4^ Targeted MS also has higher sensitivity and dynamic range than untargeted MS, measuring peptides that would otherwise be undetectable.^5^

Label-free MS-based targeted proteomic approaches such as parallel reaction monitoring (PRM)^6^ are common, but these approaches are limited by factors that include low sample throughput, error introduced by both systematic and random variation between runs, and the laborious and sometimes unsuccessful process of assay development, which involves the manual scheduling of targeted MS2 scans and may require the synthesis of peptide standards.^1^ Though recent technologies have made PRM assays resilient to chromatographic variation,^7,8^ PRM assay development remains a barrier to the wide adoption of targeted MS.

We recently reported a novel targeted proteomic method, termed GoDig, which requires neither manual PRM scheduling nor synthetic standard peptides and uses tandem mass tags (TMT) to eliminate run-to-run variability within plex and increase sample throughput up to 18-fold.^5^ Using only a data-dependent acquisition (DDA)-based spectral and elution library and a target peptide list, GoDig makes real-time decisions to automatically detect, identify, and quantify targets. A publicly available DDA-based library such as those available for >10,000 human proteins or >4,500 yeast proteins may be used,^5,9,10^ or a custom library can be assembled by fractionating a reference sample and analyzing the fractions with a collision-induced dissociation (CID)-high-resolution MS2 (HRMS2) method. With GoDig, over 2,500 abundant peptides can be identified in a single run.^5^ With the use of chromatographic priming, peptides of low abundance usually not quantifiable with untargeted methods can be quantified with a >95% success rate.^10^

Although the success rates of GoDig are suitable for most protein abundance measurements, GoDig has not yet been used to measure post-translational modifications (PTMs), which are critical to understanding biological systems and diseases; reliably quantifying single PTM-bearing peptides requires high peptide-level success rates. Here, we describe the systematic identification and reduction of several of the limitations of GoDig to result in a technology, GoDig 2.0, which is unprecedented in its convenience, flexibility, and throughput—both in terms of samples and targets. In addition to compatibility with the TMTproD 35plex reagents, which nearly double sample throughput,^11^ the success rates and target throughput of GoDig 2.0 are substantially improved by several software innovations. We show that the resulting method is suitable for quantifying PTMs and covalent compound-protein interaction sites, and showcase this by measuring differences in tau phosphorylation between brain tissues from patients with Alzheimer’s disease (AD) and age-matched controls and by profiling the dose-response profiles of reactive cysteines across biological pathways and protein classes.

## RESULTS

### GoDig 2.0 Quantifies Twice As Many Targets as GoDig 1.0 with a Challenging Target List

We used GoDigViewer^10^ to diagnose failures to quantify targets and then made six improvements upon the original GoDig implementation (“GoDig 1.0”) to produce what we call GoDig 2.0 (Fig. 1, Fig. S1). These improvements were: (1) compatibility with higher-energy collision-induced dissociation (HCD)-high-resolution MS2 (HRMS2) library data; (2) incorporation of retention time alignment into library building; (3) sequential narrow window accumulation-MS1 (MSX-SIM) monitoring; (4) dynamic target close-out; (5) the subcycle scan algorithm; and (6) TMTproD 35plex compatibility. (For extensive description and illustration of these improvements and their effects, see the Supplementary Information and Fig. S1–S8.) To benchmark GoDig 2.0, we prepared a 5-cell-line mixture 18plex (Fig. 2A) and targeted 2,000 precursors, including 1,300 precursors randomly selected from a published 4-human-cell-line fraction-based GoDig library and nearly 700 targets that were never detected over 9 priming runs and are thus not feasibly detectable in this sample (Table S1, Fig. S2–S3). As opposed to past studies in which we tested GoDig on detectable “super fliers,”^5,9^ we chose to simulate a challenging scenario in which a large target list spanned the range of abundances and ionizabilities from a fractionated dataset and was interfered with by targets that were not present or detectable in the sample. In this challenging context, GoDig 2.0 was able to quantify 911 ± 17 precursors in a single run, compared to only 401 ± 51 for GoDig 1.0—an improvement by a factor of 2.4x (Fig. 2).

**Figure 1.**
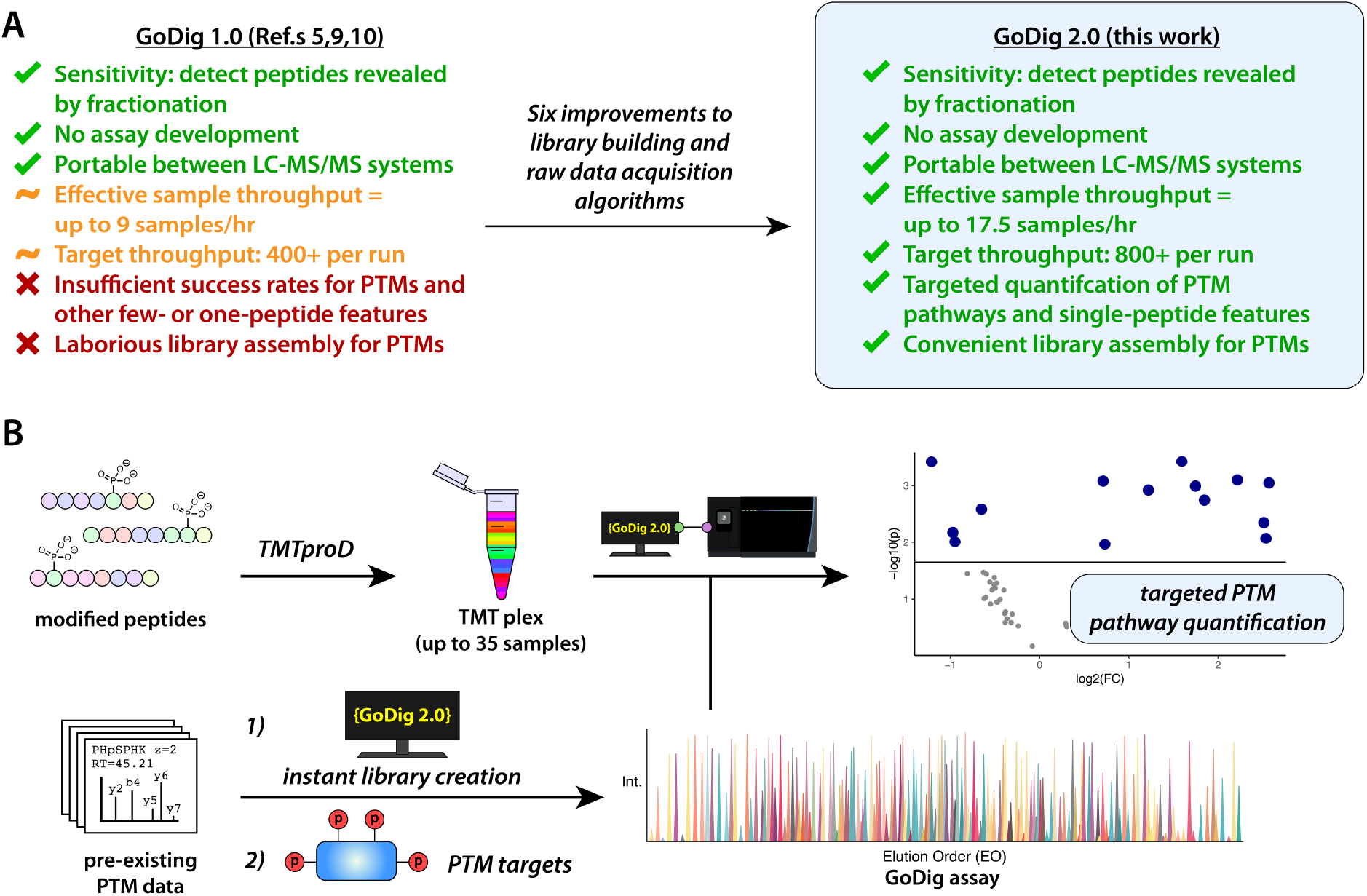
GoDig 2.0 improves on success rates, throughput, and ease of use compared to GoDig 1.0. A. Limitations and features of GoDig 1.0 and 2.0. **B**. Generic GoDig 2.0 workflow depicting the analysis of modified peptides using TMTproD 35plex reagents.

**Figure 2.**
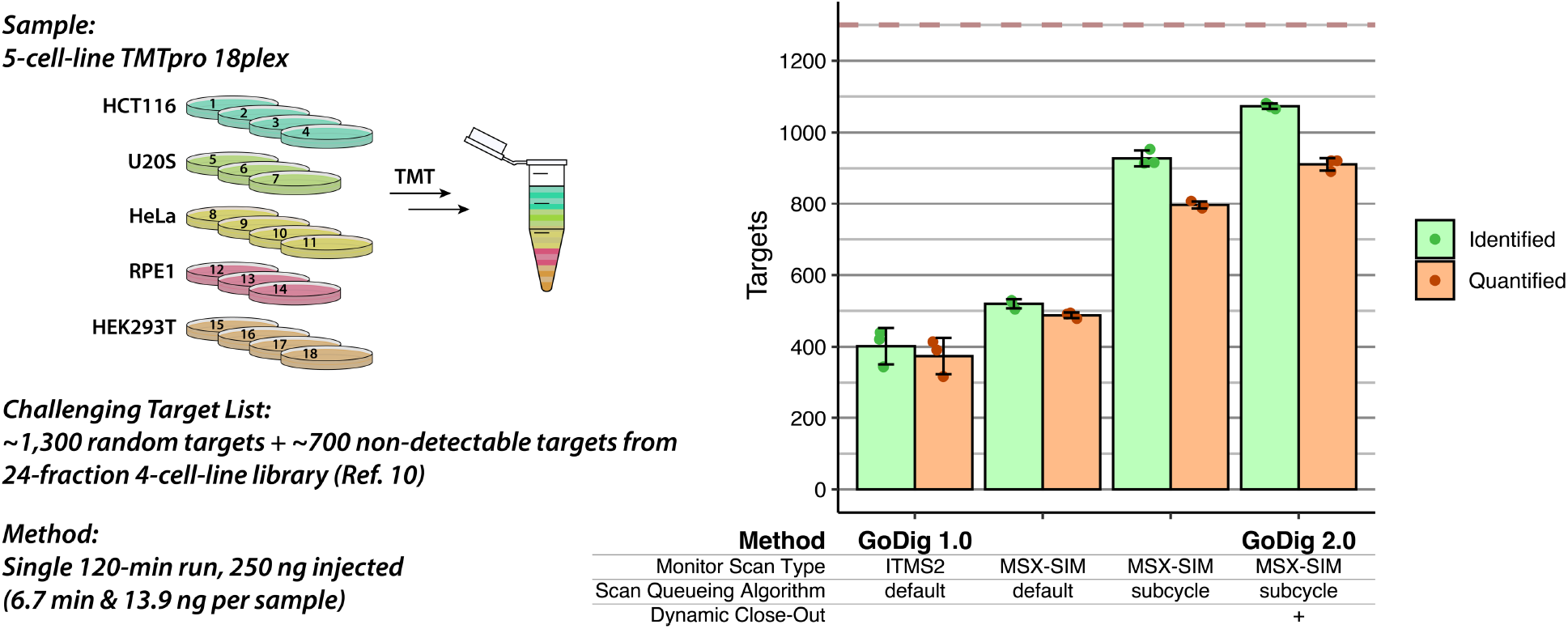
GoDig 2.0 quantifies over 2.4x more targets than GoDig 1.0 in a challenging context. For details about the target list and GoDig 2.0 features and parameters, see the Supplementary Information and Supplementary Figures. Identified = passed spectral match filters. Quantified = summed signal-to-noise exceeded 10/channel. Upper bound on performance at 1,300 targets is indicated by dashed red line. Corrected EO bins from 6 priming runs were used for all of these experiments (Ref. 10).

### GoDig 2.0 Increases Sample Throughput Via TMTproD 35plex Compatibility

To increase sample throughput and to remain current with TMT technology, we implemented features in the instrument API (iAPI) and in GoDig that make GoDig 2.0 compatible with the new TMTproD 35plex reagents.^11^ Specifically, we added “TurboTMT” (phiSDM^12^) as a feature in the iAPI and then used this to make TurboTMT a feature in GoDig 2.0, enabling the orbitrap to resolve TMTproD peaks from non-deuterated TMTpro peaks at resolution = 60k as described by Zuniga et al.^11^ To test this feature, we generated and analyzed a two-cell-line mixture sample, following the design of Zuniga et al., which (1) used all 35 TMTproD kit reagents, (2) contained N=8 samples from each cell line in both the deuterated and non-deuterated subplexes, and (3) contained three bridge channels (Fig. 3A). ^11^

**Figure 3.**
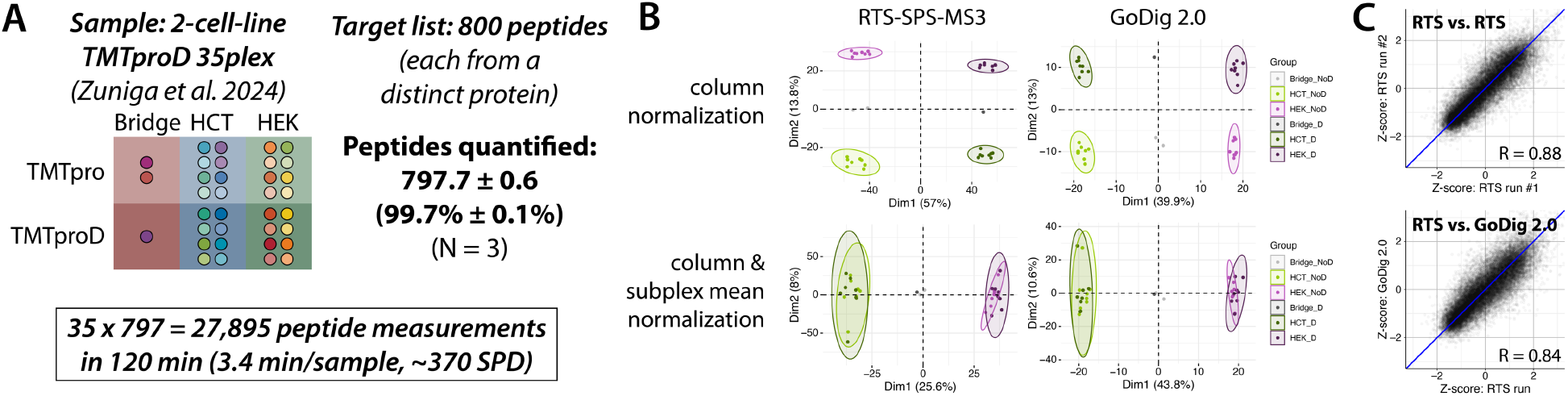
GoDig 2.0 increases effective sample throughput from 6.7 min/sample to 3.4 min/sample with TMTproD. A. TMTproD 35plex sample and target list used to show compatibility with GoDig 2.0. SPD = samples per day. **B**. Principal component analysis (PCA) using untargeted RTS-SPS-MS3 analysis with resolution = 120,000 and GoDig 2.0 with resolution = 60,000 and TurboTMT enabled. Column normalization = dividing each column (channel) by its total sum. Subplex mean normalization = dividing each measurement by the mean of measurements for that target within the subplex (i.e., all non-D channels or all D channels). **C**. Scatter plots showing the correlation of quantities between replicate untargeted RTS-SPS-MS3 runs (“RTS”) and between one RTS run and a GoDig 2.0 run. Z-scores were calculated within each MS3 scan. R = Pearson correlation coefficient.

Since GoDig 1.0 previously achieved a >95% success rate quantifying 125 proteins in the macroautophagy pathway by targeting 2 peptides per protein,^9^ we hypothesized that GoDig 2.0 could make measurements with a peptide-level success rate sufficient to quantify proteins using a single peptide per protein. We selected 800 peptides, each from a distinct protein (Table S2), and targeted each with a single peptide using GoDig 2.0 in the 2-cell-line 35plex, quantifying 797.7 ± 0.6 (99.7%) of the peptides in a single run. TurboTMT enabled the PCA analysis to resemble that of an untargeted RTS-synchronous precursor selection (SPS)-MS3 analysis (Fig. 3B). GoDig 2.0 requires 3.4 min per sample while simultaneously taking advantage of 120 min of chromatographic separation, something not possible with label-free methods; allowing 15 min for sample loading, this corresponds to ~370 samples per day.

### Streamlined Targeted Pathway Phosphoproteomics Is Enabled by Data Repurposing in GoDig 2.0

Phosphorylation is a common PTM vital to many biological processes and diseases, including cancer and Alzheimer’s disease (AD).^13,14,15^ Targeted MS has the ability to simultaneously quantify phosphorylation sites spanning a whole signaling pathway.^16,17,18^ We sought to apply GoDig 2.0 to targeted phosphoproteomics, but the assembly of a broadly useful library such as those used for protein abundance measurements^5,9^ was not feasible because the presences and abundances of phosphoproteins are strongly dependent on the biological context. Therefore, we envisioned a “data repurposing” approach, where previously acquired untargeted data are compiled into libraries (Fig. 4A). We implemented HCD fragmentation as an option for IDMS2 scans, making GoDig compatible with high-field asymmetric-waveform ion mobility (FAIMS)-HCD-HRMS2 data, the typical data format for untargeted phosphoproteomic TMT studies.^19,20^ Second, we incorporated retention time (RT) alignment into library building, allowing the integration of previously acquired datasets that were acquired under different chromatographic conditions (Fig. S8).

**Figure 4.**
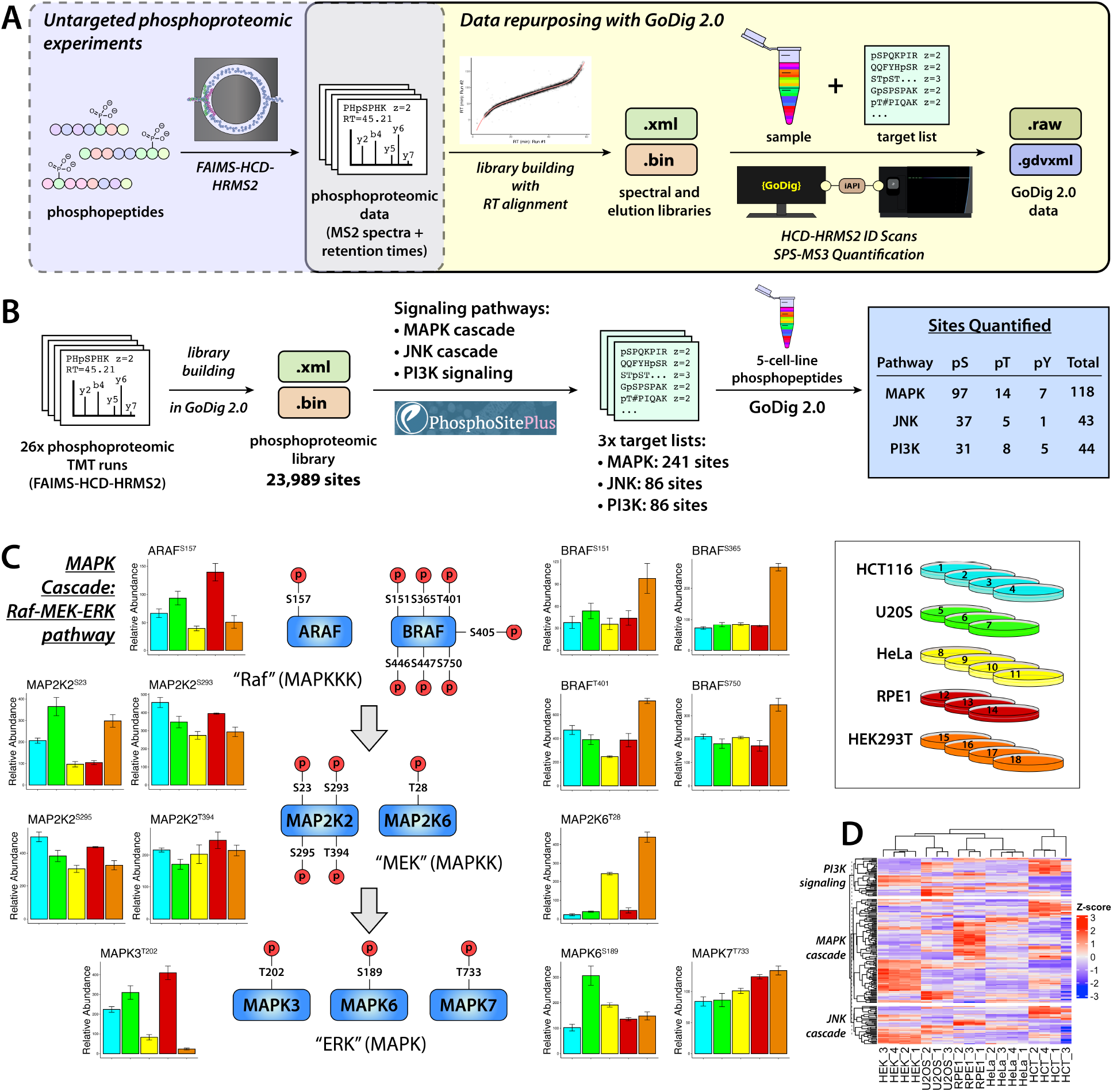
GoDig 2.0 enables the repurposing of existing phosphoproteomic data for streamlined targeted pathway analysis. A. Data repurposing workflow depicted with phosphoproteomic data as an example. Previously acquired untargeted LC-MS/MS data, including PTM analyses, regardless of fragmentation mode, buffer gradient, or ion mobility-based separation, can be built into spectral and elution libraries using GoDig 2.0. **B**. A three-pathway targeted phosphoproteomic experiment performed with GoDig 2.0. For each pathway, 3 priming runs and 1 analytical run were performed (Ref. 10). **C**. Examples of phosphoprotein abundance differences in the Raf-MEK-ERK portion of the MAPK cascade in a 5-cell-line 18plex. **D**. Heat map of Z-transformed GoDig 2.0 measurements. Rows are phosphoproteins and columns are cell line replicates.

Using 26 previously acquired phosphoproteomic runs with varying sample composition and chromatography, we built a library containing 36,550 peptides representing 23,989 different phosphorylated forms of 5,860 human proteins (Fig. 4B). We retrieved phosphoproteins in the MAPK cascade, JNK cascade, and PI3K signaling cascade from PhosphoSitePlus, then enriched phosphopeptides from the 5-cell-line TMTpro 18plex described above and targeted each of these pathways using three priming runs and one analytical run.^10^ Despite the differences between these runs and samples and the runs and samples used for the library, we successfully quantified 118, 43, and 44 phosphoproteins each with GoDig 2.0 (Fig. 4B, Table S3).

The mitogen-activated protein kinase (MAPK) cascade is a well-known pathway in development and cancer, facilitating cellular growth and proliferation.^21,22^ A nexus of this pathway is the Raf-MEK-ERK cascade, whose sites are regulated at the pathway level (e.g., most of these sites are downregulated in HeLa cells), but also at the homolog and site levels (Fig. 4C). The heat map in Fig. 4D shows at site-specific resolution the vast differences between cell lines in how they utilize these signaling pathways.

### A GoDig 2.0-Based Screen-To-Validation Workflow Measures Hyperphosphorylated Tau in Postmortem Tissue from Human Patients with Alzheimer’s Disease

Having demonstrated the utility of GoDig 2.0 in targeted phosphoproteomics, we sought to apply it to a pilot study of Alzheimer’s disease (AD), for which tau hyperphosphorylation is a known hallmark and pathogenic process.^14^ We acquired postmortem human temporal cortex samples from N = 10 patients with AD and N = 8 control patients with no cognitive impairment (NCI), selected to minimize effects of age and sex (Fig. 5A, Table S4). To maximize proteomic depth, we analyzed the 18plex in four injections using an HCD-HRMS2 method without FAIMS; although MS2-level analysis without FAIMS is notoriously quantitatively inaccurate,^19,20,^23 we planned to validate any observed changes using quantitatively accurate MS3-based GoDig 2.0 (Fig. 5A).

**Figure 5.**
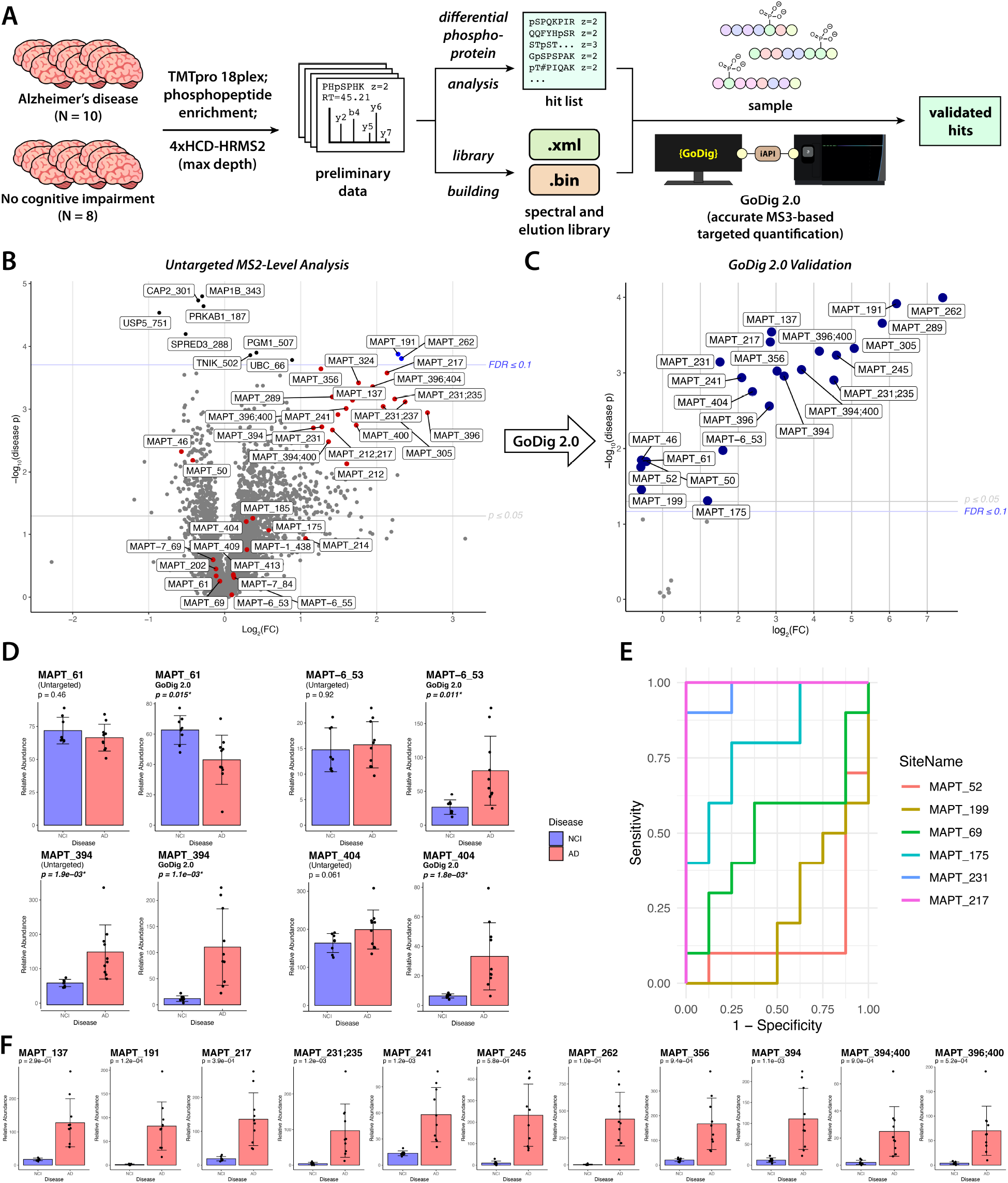
Targeted Phosphoproteomics in Brain Tissue from Patients with Alzheimer’s Disease using GoDig 2.0. A. Screen-to-validation workflow. **B**. Volcano plot showing results of untargeted HCD-HRMS2 analysis with significant and Tau (MAPT) phosphoproteins highlighted. Semicolon indicates that the protein is phosphorylated at two sites. FDR threshold was set using the Benjamini-Hochberg threshold correction method (Ref. 24). **C**. Volcano plot showing results of GoDig analysis of Tau sites. **D**. Quantification of a selection of sites with untargeted HCD-HRMS2 and GoDig 2.0. **E**. Receiver operating characteristic (ROC) plots of a selection of Tau phosphorylation sites. **F**. Quantitative data from the 11 Tau sites with perfect ROC curves (area under the curve = 1). MAPT represents isoform 8 of tau on Uniprot (P10636-8), and MAPT-1, MAPT-6, and MAPT-7 represent isoforms 1, 6, and 7, respectively.

Of the 5,695 phosphoproteins quantified by untargeted HCD-HRMS2, only 10 were statistically significant at FDR ≤ 0.1 (estimated by the Benjamini-Hochberg [BH] method^24^) (Fig. 5B, Table S5). However, pTau191 (MAPT_191) and pTau262 (MAPT_262) had the strongest significant changes, exceeding a 4-fold increase in AD, and additional tau phosphopeptides had strong fold changes below the FDR threshold. We used GoDig 2.0 to quantify 35 tau phosphopeptides representing 30 forms of phosphorylated tau (Table S6), using the untargeted HCD-HRMS2 data as the library (possible only with GoDig 2.0). Strikingly, at FDR ≤ 0.1, GoDig 2.0 revealed 23 significantly altered pTau forms, including 4 that were decreased in AD (Fig. 5C). Fig. 5D shows four pTau forms for which the increased accuracy of GoDig 2.0 revealed a fold change that was diminished when measured with untargeted MS, including three whose *p*-values were nonsignificant when measured with untargeted MS. Thus, the validation step of the GoDig 2.0-based screen-to-validation workflow can reveal differential analytes both by reducing the sensitivity cost of the BH method^25^ and by improving quantitative accuracy.

In this cohort, some pTau forms discriminate between patients with AD and those with NCI. Fig. 5E shows receiver operating characteristic (ROC) plots for a selection of pTau forms, including pTau217, a well-known biomarker of AD that has been used successfully in blood plasma;^26^ with GoDig 2.0, pTau217 discriminated perfectly between disease states, giving an area under the curve (AUC) of 1. We found 10 additional pTau forms that achieved this same feat (Fig. 5F); future work will investigate whether this ensemble of hyperphosphorylated tau forms can improve on pTau217 as a biomarker for AD.

### GoDig 2.0 Reveals Cell-Line-Specific Polyubiquitin Branching

To further validate the utility of GoDig 2.0 in studying PTMs, we sought to quantify ubiquitylation sites.^27^ One important subclass is polyubiquitylation (polyUb), in which ubiquitin is ubiquitylated at one or more of 7 lysines: K6, K11, K27, K29, K33, K48, and K63.^27^ We searched for the Lys-diGly (KGG) modification (produced by trypsinization of ubiquitylated proteins) in the published 4-cell-line-based GoDig library dataset (Fig. S9A).^10^ The library contains 216 KGG sites, including 6 out of the 7 possible modified KGG sites on ubiquitin (Fig. S9B). In our 5-cell-line 18plex, we quantified 73 KGG-bearing peptides, including peptides covering all 6 polyUb branching sites. The analysis revealed differences in polyUb branching between cell lines; this assay may help reveal the functions of these sites, some of which are poorly understood^27^ (Fig. S9C).

### GoDig 2.0 Enables High-Throughput Quantification of Electrophilic Compound-Protein Interaction Sites

Having applied GoDig 2.0 to endogenous PTMs, we sought to target a non-natural modification useful for drug discovery: desthiobiotin (Fig. 6A). The electrophilic desthiobiotin-iodoacetamide (DBIA) probe has been used to quantify compound-protein interactions at cysteine site-level resolution by competing with exogenous electrophilic compounds for cysteine engagement; this activity-based protein profiling (ABPP) approach has been coupled with TMT for a streamlined cysteine ABPP (SLC-ABPP) workflow which can be performed in a 96-well-plate format.^28,29^ In order to use GoDig 2.0 for targeted SLC-ABPP, we built a library of 20,946 cysteines by enriching and fractionating DBIA-bound peptides from a large-scale K562 cell digest (Fig. 6A). We then treated native lysates from K562 cells with serial dilutions of each compound alongside a DMSO-treated negative control, with N=2 per condition (except for N=1 for the 500-µM condition in the deuterated 17-channel subplex) (Fig. 6B).

**Figure 6.**
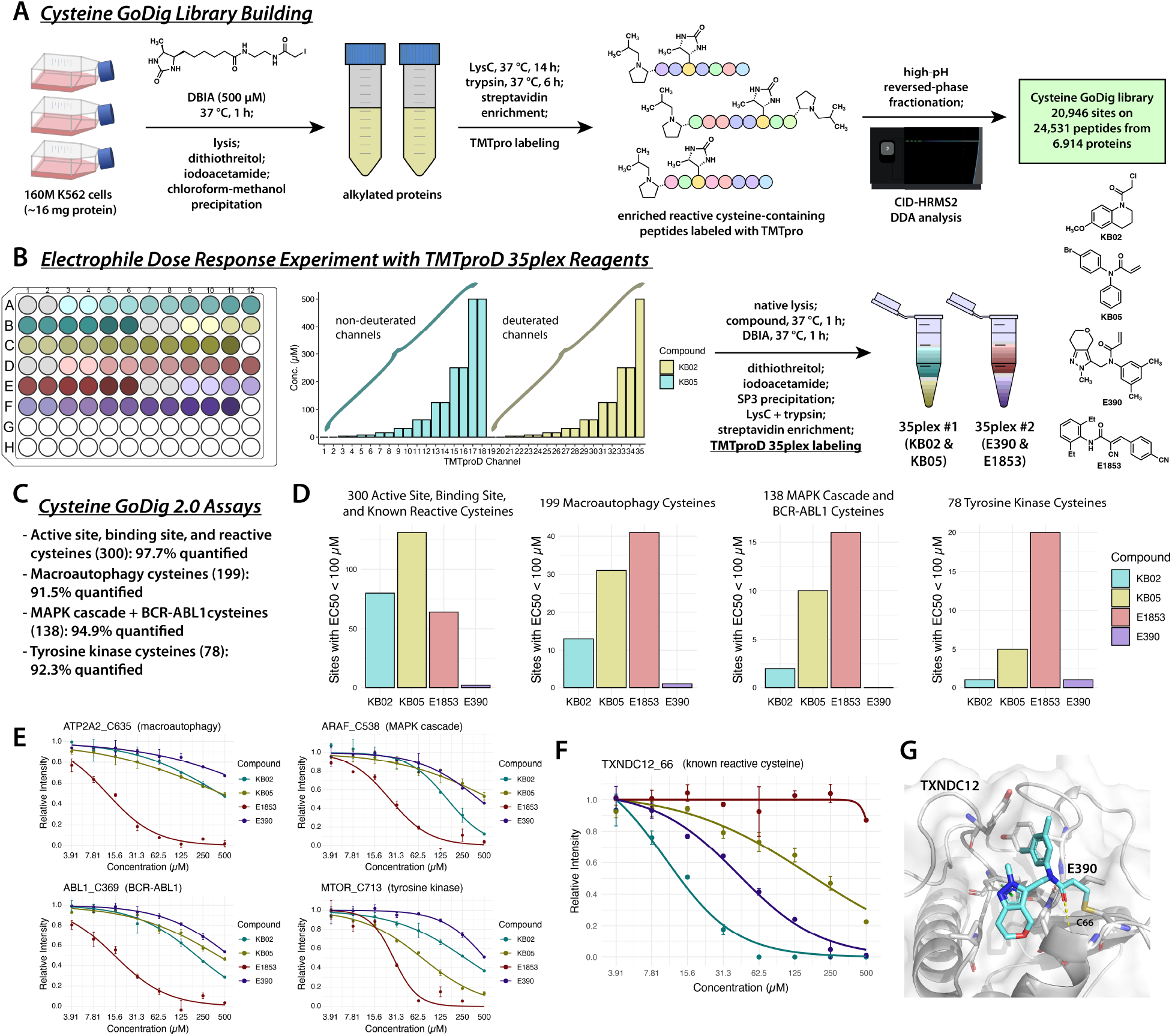
Targeted Streamlined Cysteine Activity-Based Protein Profiling (SLC-ABPP) with GoDig 2.0. A. Preparation of a GoDig library consisting of 20,946 cysteines that are reactive in live K562 cells. **B**. Dose-response experiment workflow. Gray-colored wells were treated with DMSO as a negative control. **C**. List of targeted pathway and protein class assays. Number in parentheses is the number of cysteines in the target list. **D**. Hit counts from the four assays. **E**. Dose-response curves from different pathways exemplifying the superior potency of E1853 over KB02 and KB05. **F**. Dose-response curves for TXNDC12^C66^. **G**. Energy-minimized structure of E390 bound to C66 on TXNDC12.

We targeted four different sets of cysteines in the two resulting 35plexes: first, we targeted sites that are annotated as “active site” or “binding site” cysteines on Uniprot.org, alongside sites that are known to be reactive to at least one of the electrophilic scout fragments KB02 and KB05 (as positive controls), totaling 300 sites;^30,31^ next, we targeted 199 cysteines on proteins in the macroautophagy pathway and 138 in the MAPK cascade using the QuickGO web tool as described previously;^10^ and finally, we targeted 78 sites on tyrosine kinases, a class of proteins that are important participants in intracellular signaling and are relevant to various cancers (Fig. 6C, Table S7).^32^ In a single injection, the quantification success rates of the resulting GoDig assays ranged from 91–97%. In the first target list, more sites reacted with KB02 and KB05 than E1853 or E390, but in the others, which were not biased by prior knowledge of reactivity, E1853 reacted with the most cysteines (Fig. 6D–E, Table S8–9); we attribute this reactivity to the electron-withdrawing character of the two nitrile groups as well as the hydrophobicity and planarity of the compound, which may facilitate binding to protein surfaces. Strikingly, throughout all four target lists, only 5 sites reacted with E390 with an EC_50_ of <100 µM, including cysteine 66 on TXNDC12, an endoplasmic reticulum thioreductase known to inhibit lipid peroxidation and ferroptosis.^33^ E390 reacted more strongly with TXNDC12^C66^ than KB05, reversing the overall reactivity trend (Fig. 6F). To understand this selectivity, we performed molecular docking in Maestro (Schrödinger) of E390 with the AlphaFold 3.0 model of TXNDC12 (Fig. 6G). E390 is predicted to interact with a surface-exposed hydrophobic region of TXNDC12, wherein the additional flexibility afforded by the methylene spacer of the (pyrano[4,3-c]pyrazolyl)methyl substituent allows for hydrophobic contacts with the protein and the acquisition of a π-π interaction with Phe140. The targeted generation of hundreds of dose-response curves across four pathways and protein classes, at a throughput of 3.4 min per replicate, showcases the power of GoDig to characterize potential lead compounds across pathways of therapeutic interest.

## DISCUSSION

Here, we have shown that GoDig 2.0 gives significantly higher success rates than its already widely capable predecessor, GoDig 1.0. We achieved this primarily by increasing time efficiency and reducing scan delays, using three innovations new to targeted MS: MSX-SIM monitoring, the subcycle scan queueing algorithm, and dynamic close-out. These enable multiplexed targeted studies of specific sites, as demonstrated here with post-translationally and chemically modified peptides. Beyond phosphorylation, ubiquitylation, and DBIA-modified cysteines, this approach should be applicable to any other PTM or synthetic modification amenable to MS. We have also increased sample throughput by making GoDig compatible with TMTproD reagents, which decrease instrument time to as little as 1.7 minutes per sample per gradient hour; allowing 135 min per 35plex, this equates to >370 samples per day, which could be further increased by shortening the gradient if allowed by the target list.

To apply these improvements to targeted PTM quantification, we implemented a “repurposing” approach which rapidly builds GoDig libraries from existing datasets, regardless of chromatographic variation or fragmentation mode. This repurposing approach lends itself naturally to a screen-to-validation approach wherein the same sample is first analyzed with untargeted MS to identify statistically significant hits and then with GoDig 2.0 for rapid validation with higher quantitative accuracy. We performed this workflow on a small study of human AD, recapitulating a known phosphoproteomic biomarker, pTau217, and revealing 10 more phosphorylation sites that could be pursued for inclusion in future diagnostic assays. In addition, we generated a GoDig library of 23,989 phosphorylation sites that is now publicly available via the PRIDE^34^ repository (PXD065227).

High throughput is desirable in drug discovery both in terms of samples and sites, since often the therapeutic target is not a single molecule, but a phenotype that arises from a biological pathway consisting of several proteins. We demonstrated the ability of GoDig 2.0 to target tens to hundreds of cysteines across whole biological pathways or protein classes at quantification success rates of 91–97%. We generated a library of 20,946 cysteines (also available in the PRIDE repository) and developed assays that are usable by any researcher with an Orbitrap Eclipse or Ascend via the GoDig 2.0 software, which is freely available at https://gygi.hms.harvard.edu/. Future work will expand on these target lists, increase the depth of publicly available GoDig libraries, and broaden the libraries to additional cell lines and biological contexts. Together, these results show that GoDig 2.0 is suitable for quantification of PTMs, synthetic modifications, and proteins represented by single peptides, with unprecedented sample throughput.

## ONLINE METHODS

### Cell Culture and Compound Treatment

For the 5-cell-line mixture and for the 2-cell-line TMTproD 35plex, cells were purchased from American Type Culture Collection (HCT116: #CCL-247; U2OS: #HTB-96; HeLa: #CRM-CCL-2; RPE1: #CRL-4000; HEK293T: #CRL-3216), thawed according to the manufacturer’s protocol, and grown on 15-cm plates in Dulbecco’s modified eagle medium (DMEM) with 10% fetal bovine serum (FBS). Cells were harvested on ice by removing media and washing 1x with ice-cold phosphate-buffered saline (PBS). Cells were then scraped into 1 ml of ice-cold PBS and pelleted by centrifugation. After removing the supernatant, cells were frozen in liquid nitrogen and stored at –80 °C.

For the reactive cysteine GoDig library, K562 cells (ATCC: #CCL-243) were grown in sterile polystyrene T-75 culture flasks in 20 ml Roswell Park Memorial Institute (RPMI)-1640 media with 10% FBS and glutamine. When 160 million cells were present in 8 culture flasks, cells were centrifuged at 300 rcf for 5 min at 4 °C and the supernatants were removed. Cells were resuspended in the same media at 3 million cells/ml and treated with 500 µM DBIA for 1 hour @ 37 °C, shaking every 10–15 min. Cells were then centrifuged at 300 rcf for 5 min at 4 °C, resuspended in 20 ml ice-cold PBS, centrifuged and resuspended in ice-cold PBS a second time to wash, combined into two separate suspensions of 80M cells each, and then centrifuged and supernatant removed. The two resulting cell pellets were frozen in liquid nitrogen and stored at –80 °C.

For the covalent compound dose-response experiment, K562 cells were grown in T-75 flasks to near-confluence, collected by centrifugation, washed twice with ice-cold PBS, then frozen in liquid nitrogen and stored at –80 °C as described above. Frozen cells were then resuspended and lysed in native lysis buffer (PBS, pH = 7.4 + 0.1% NP-40) by extrusion through a 21-gauge needle. The crude lysate was then subjected to EpiShear™Probe Sonicator (Active Motif) for further homogenization (5 min, 3 s on, 3 s off, 50% amplitude) on ice. The resulting lysate was centrifuged at 1,400 rcf for 5 min. and the supernatant containing the soluble proteome was collected. The protein concentration was measured by BCA assay and diluted to 2 µg/µl with lysis buffer. 15 µl of lysate containing 30 µg of protein was then loaded into each of 35 wells in a 96-well plate. 5 µl of compound solution in lysis buffer was added to a final concentration between 0– 500 µM (in each experiment, serial 2x dilutions were prepared from 500 µM to 3.90625 µM and a 0-µM control condition was included) and incubated for 1 h. 5 µl of DBIA solution in lysis buffer was then added to a final concentration of 500 µM and incubated in the dark for 1 h.

### Postmortem Brain Tissue Acquisition and Homogenization

Post-mortem fresh-frozen temporal cortex tissue samples were obtained from the Stanford/VA/NIA Aging Clinical Research Center (ACRC) with approval from local ethics committees and patient consent. Patient characteristics are presented in Table S4. AD diagnosis is based on pathology whereas patients with NCI had no clinical dementia or pathological AD diagnosis. All procedures were carried out on ice in a 4 °C cold room as rapidly as possible. 1000 mg of brain tissue was thawed on ice in 800 µl 200 mM EPPS, pH = 8.5 + 8 M urea with Pierce protease and phosphatase inhibitors (Thermo Fisher Scientific) and then homogenized with a VWR 200 homogenizer using 4x 5-s pulses spaced 3 s apart. The homogenates were then centrifuged at 1,200 rcf for 10 min at 4 °C and the supernatant was collected. The resulting homogenate was then stored at –80 °C.

### Sample Preparation

For the 5-cell-line mixture, pellets were thawed and resuspended in lysis buffer (200 mM EPPS, pH = 8.5 + 8 M urea + Pierce protease and phosphatase inhibitors) and ruptured with probe sonication (5 s at 4 °C and 50% amplitude) and then sheared by 10x extrusion through a 21-gauge, 1.25”-length needle. Protein concentration was measured with BCA, 1 mg protein per lysate was aliquoted, and volume was brought to 1 ml with lysis buffer. Lysates were treated with 5 mM dithiothreitol (DTT) for 30 min at 37 °C, 10 mM iodoacetamide (IAA) in the dark for 30 min at room temperature, and then 10 mM DTT for 15 min at room temperature. Each reduced and alkylated lysate was then transferred to a 15-ml tube and 4 ml methanol, 1 ml chloroform, and 3 ml water were added. The mixture was vortexed thoroughly and inverted 5x, then centrifuged for 15 min at 4,696 rcf at room temperature. The aqueous and organic layers were then both removed, leaving a protein precipitate, which was then washed with 4 ml methanol by vortexing and then centrifuging at 4,696 rcf for 30 min at room temperature. The supernatant was then removed and each proteome was resuspended in 1 ml 200 mM EPPS, pH = 8.5 by vortexing and bath sonication. The proteins were then digested with LysC (Wako/Fujifilm) at 1:100 E:S ratio at 37 °C overnight while shaking, and then further digested with trypsin (Thermo Scientific) at 1:100 E:S ratio at 37 °C for 6 h. The peptides were then acidified by adding 100 µl formic acid and then cleaned up on a 200-mg Sep Pak column (Waters) according to the manufacturer’s protocol. The elution buffer was then removed in a vacuum centrifuge. Each 1-mg digest was then subjected to phosphopeptide enrichment using a High-Select™ Fe-NTA Phosphopeptide Enrichment Kit (Thermo Scientific) according to the manufacturer’s protocol, retaining all “flow-thru” (non-phosphorylated/non-enriched) peptides. All peptides were then dried in a vacuum centrifuge. Flow-thru peptides were cleaned up on a 200-mg Sep Pak column (Waters) according to the manufacturer’s protocol and phosphopeptides were resuspended in 100 µl 10% formic acid and cleaned up using in-house StageTips (Rappsilber, Mann & Ishihama, *Nature Protocols* **2007**, *2*, 1896), and then all peptides were dried in a vacuum centrifuge. Each flow-thru peptide sample was resuspended in 200 µl 1.5% acetonitrile + 0.5% formic acid in water and then 20 µl (~100 µg) was aliquoted for TMT labeling and dried in a vacuum centrifuge. Each sample (~100 µg flow-thru, ~5–40 µg phosphopeptide) was then resuspended in 100 µl 200 mM EPPS, pH = 8.5, then 30 µl acetonitrile was added, then 20 µl of a 12.5 µg/µl solution of a different TMTpro reagent was added to each biological sample. The reactions were shaken at 1,000 rpm for 1 h at room temperature. For ratio check, to 150 µl 2% formic acid in water was added 2 µl of each reaction (this was performed separately for the 18 flow-thru samples and the 18 phosphopeptide samples) and then all reactions were frozen at –20 °C. These ratio-check 18plexes were cleaned up using StageTips and analyzed by HCD-HRMS2 (see below) (60-min gradient, 1 µg injection) on an Orbitrap Eclipse or Exploris 240 mass spectrometer (Thermo Fisher Scientific). The reactions were then thawed, 10 µl 5% hydroxylamine was added, and the reactions were incubated for 15 min at room temperature to quench. >50% of the volume was evaporated with a vacuum centrifuged, 90 µl 10% formic acid + 3% acetonitrile was added to each sample, and the phosphopeptides were combined according to the ratio check measurements. The individual non-phosphopeptide samples/channels and the phosphopeptide 18plex were cleaned up on 50-mg Sep Pak columns (Waters) according to the manufacturer’s protocol and then dried in a vacuum centrifuge. The phosphopeptide 18plex was resuspended in 100 µl 3% acetonitrile + 1% formic acid for mass spectrometric analysis. Each non-phosphopeptide sample/channel was resuspended in 100 µl 3% acetonitrile + 1% formic acid in water and then the samples/channels were combined according to the ratio check measurements.

For the 2-cell-line TMTproD 35plex, clean digests were prepared in the same manner described above for the 5-cell-line 18plex (stopping short of phosphopeptide enrichment). Each of the 16 peptide samples was resuspended at 1 µg/µl in 200 mM EPPS, pH = 8.5, and then from each was aliquoted 2x 15-µl (15-µg) replicates (one to be labeled with a non-deuterated TMTpro reagent, the other to be labeled with a deuterated TMTproD reagent) and each of those replicates was diluted 2x with 15 µl 200 mM EPPS, pH = 8.5. To a single tube was added 4 µl from each of the original 16 peptide samples, resulting in a 64-µl bridge sample, which was then also diluted 2x to 0.5 µg/µl and then aliquoted into 3x 30-µl (15-µg) replicates (two to be labeled with TMTpro-126 and TMTpro-127N, the third to be labeled with TMTpro-127D). To each of the resulting 35 30-µl samples was added 6 µl acetonitrile and 7 µl of a TMTproD reagent (12.5 µg/µl in acetonitrile), with the following channel layout: 1–2 = Non-D Bridge; 3–10 = biologically distinct HEK293T samples; 11–18 = biologically distinct HCT116 samples; 19 = D Bridge; 20–27 = replicates of 3–10; 28–35 = replicates of 11–18. The reactions were incubated, ratio-checked, and quenched as described above. The hydroxylamine-quenched peptides were combined according to the ratio check measurements, >30% of the volume was evaporated in a vacuum centrifuge, and the 35plex was acidified to pH < 3 with formic acid and then cleaned up on a 100-mg Sep Pak (Waters) according to the manufacturer’s protocol. The 35plex was then dried in a vacuum centrifuge and resuspended in 3% acetonitrile + 1% formic acid in water for LC-MS/MS analysis.

For the human brain samples, the homogenates were thawed on ice, diluted 2x with ice-cold lysis buffer (200 mM EPPS pH = 8.5 + 8 M urea + Pierce phosphatase and protease inhibitors), probe sonicated 3x for 5 s at 30% amplitude at 4 °C, and centrifuged at 15,000 rcf for 5 min at 4 °C, and the supernatant was collected. The protein concentration was measured via BCA assay. 100 µg protein per homogenate was aliquoted and all volumes were increased to 100 µl with lysis buffer. The lysates were reduced with DTT, alkylated with IAA, and quenched with DTT as described above, then 400 µl methanol, 100 µl chloroform, and 300 µl water were added to reach a total volume of 900 µl. The biphasic suspensions were vortexed thoroughly and then centrifuged at 17,000 rcf at room temperature. The aqueous and organic layers were removed by aspiration and the protein precipitate was washed in 400 µl methanol by vortexing and then centrifuging for 30 min at 17,000 rcf at room temperature. The supernatant was removed and the proteins were resuspended in 100 µl 200 mM EPPS, pH = 8.5. LysC was then added at a 1:100 E:S ratio and incubated at 37 °C overnight with shaking, then trypsin was added at 1:100 E:S ratio and incubated at 37 °C for 6 h with shaking. 30 µl acetonitrile was then added, then TMTpro reactions were carried out by adding 20 µl of a 12.5 µg/µl solution of each TMTpro reagent and then incubating for 1 h at room temperature while shaking at 1,000 rpm. The reactions were ratio-checked, quenched, combined, acidified, and cleaned up on a 200-mg Sep Pak as described above. Phosphopeptides were enriched from the resulting 1.8-mg 18plex using the High-Select™ Fe-NTA Phosphopeptide Enrichment Kit (Thermo) according to the manufacturer’s protocol. The eluted phosphopeptides were dried in a vacuum centrifuge, resuspended in 100 µl 3% acetonitrile + 1% formic acid, cleaned up using a StageTip as described above, dried in a vacuum centrifuge, and resuspended in 20 µl 3% ACN + 1% FA for HCD-HRMS2 analysis (see below) and GoDig 2.0 analysis (see below).

For the reactive cysteine GoDig library, each of the two 80-M (~8 mg) cell pellets was thawed in 5.3 ml lysis buffer and lysed by vortexing then by passing 10x through a 21-gauge, 1.25”-length needle. The lysates were then reduced with DTT, alkylated with IAA, quenched with DTT, precipitated in chloroform + methanol + water, washed, and digested as described above. The digests were then cleaned up on a 2-g Sep Pak (Waters) according to the manufacturer’s protocol. The digests were then aliquoted into 32x 500-µg digests which were all subjected to the same enrichment procedure. Each 500-µg digest was resuspended in 1 ml Dulbecco’s PBS (DPBS) and 100 µl of magnetic streptavidin beads (Thermo Fisher Scientific, cat. # PI88817) was added and the mixtures were rotated overnight at room temperature. Tubes were placed on a magnetic stand for ≥2 min and the supernatant was decanted. The beads were washed with 3x 1 ml DPBS, then 1 ml 0.1% sodium dodecyl sulfate in DPBS, then 3x 1 ml water before eluting with 2x 500 µl 50% + 1% FA, combining all eluates into the same tube which was then frozen in liquid nitrogen and freeze-dried in a lyophilizer. Enriched peptides were then quantified using the Pierce Quantitative Fluorometric Peptide Assay (Thermo Scientific) according to the manufacturer’s protocol, resuspended at 1 µg/µl, and labeled with surplus TMTpro reagents as described above before quenching, removing acetonitrile in a vacuum centrifuge, and acidifying as described above. The enriched peptides were then fractionated using the Pierce High pH Fractionation Kit (Thermo Fisher Scientific) as described in the manufacturer’s protocol. The fractions were dried in a vacuum centrifuge, resuspended in acidic buffer, cleaned up with StageTip as described above, and then analyzed by untargeted CID-HRMS2 to generate the GoDig library.

For K562 cells treated with electrophilic compounds, 4 µl of SP3 beads (1:1 mixture of hydrophobic and hydrophilic type, 50 mg/ml, cat. # 45152105050250 and cat. # 65152105050250) were added to lysates that had been incubated with DBIA for 1 h, followed by addition of 40 µl absolute ethanol supplemented with 20 mM DTT. This mixture was incubated for 10 min with mild shaking before placing he plate on a magnetic stand to aspirate the supernatant. Beads were washed once with 150 µl of 80% ethanol and resuspended in 30 µl of lysis buffer supplemented with 20 mM IAA and incubated in the dark for 30 min with vigorous shaking. 60 µl of absolute ethanol supplemented with 20 mM DTT was added and mild shaking was performed before three washes with 150 µl of 80% ethanol were performed. The supernatant was removed and 30 µl of 200 mM EPPS, pH = 8.5 containing 0.3 µg LysC (Wako/Fujifilm) was added. After a 3-h incubation at room temperature, 5 µl of 200 mM EPPS, pH = 8.5 containing 0.3 µg trypsin was added and incubated with the beads at 37 °C overnight. To the digest (with beads) were added 9.5 µl of acetonitrile and 6.5 µl of a 10 µg/µl solution of the corresponding TMTproD 35-plex reagent, followed by gentle mixing for 1 h at room temperature. The reactions were quenched by adding 7 µl of 5% hydroxylamine. All samples were combined, rinsing the wells twice with 100 µl of 1% formic acid to gather all peptides, and the combined sample was dried in a vacuum centrifuge. The resulting 35plex was resuspended in 10% formic acid in water and the supernatant was separated from the beads using a magnetic stand. The supernatant was then desalted using a 100-mg Sep Pak (Waters) according to the manufacturer’s protocol and then dried in a vacuum centrifuge. The peptides were then resuspended in 460 µl of 100 mM HEPES buffer, pH = 7.4, and then 80 µl of pre-washed Pierce™ High-Capacity Streptavidin Agarose beads (cat. # 20359) was added. This mixture was incubated at room temperature with gentle agitation for 2–4 h before being loaded into an Ultrafree-MC centrifugal filter (hydrophilic PTFE, 0.22 µm pore size) and centrifuged at 800 rcf for 30 s. Beads were washed twice with 300 µl of 100 mM HEPES, pH = 7.4 + 0.05% NP40, then three times with 350 µl of 100 mM HEPES, pH = 7.4, and then once with 400 µl of water. Peptides were then eluted sequentially by (1) incubation with elution buffer (80% acetonitrile + 0.1% formic acid in water) for 20 min at room temperature; (2) incubation with elution buffer for 10 min at room temperature; and (3) incubation with elution buffer for 10 min at 72 °C. The combined eluent was dried in a vacuum centrifuge and cleaned up with a StageTip prior to LC-MS/MS analysis (see below).

### Liquid Chromatography-Tandem Mass Spectrometry

All LC-MS/MS experiments were performed on an Orbitrap Eclipse mass spectrometer (Thermo Fisher Scientific) equipped with an EASY-nLC 1200 system (Thermo Fisher Scientific) bearing a 30-cm column packed with ReproSil-Pur C18-coated beads that are 2.4 µm in diameter (Dr. Maisch GmbH, part no. r124.aq.0001) according to the FlashPack protocol (Kovalchuk, Jensen, and Rogowska-Wrzesinska, *Mol. Cell. Proteomics* **2019**, *18*, 383) and heated to 60 °C. The column was mounted onto a nanospray ion source designed and constructed in-house and operated in positive mode at 2600 V with the transfer tube heated to 300 °C. All spectra were acquired in centroid mode with the RF Lens set to 30% amplitude. Unless otherwise specified, all LC-MS/MS experiments were performed with the following chromatographic parameters. An estimated 1 µg of peptide was injected and separated using the following 2-h gradient at 500 nl/min: a linear increase from 5% Buffer B (5% water + 0.125% formic acid in acetonitrile) in Buffer A (5% acetonitrile + 0.125% formic acid in water) to 28% Buffer B over 97 min, followed by a linear increase to 44% Buffer B over 13 min, followed by a linear increase to 100% Buffer B over 5 min, followed by isocratic flow at 100% Buffer B for 5 min.

Unless otherwise specified, GoDig was performed without priming runs with the following parameters. A bin width of 0.5 min was used and a monitor window of ±3 bins was used so that, on average, target monitoring began 1.5 min before the predicted elution and ended 1.5 min afterwards. For GoDig 1.0 experiments, a monitor dynamic exclusion duration of 5 s was used, whereas for experiments using the subcycle system, a monitor dynamic exclusion duration of 0 s was used. For GoDig experiments using the subcycle system with ion trap MS2 (ITMS2) monitoring, a subcycle size limit of 5 targets was used, whereas for experiments using the subcycle system with MSX-SIM monitoring, a subcycle size limit of 15 targets was used and up to 5 targets were monitored per MSX-SIM scan. For ITMS2 monitoring, an isolation width of 0.5 Th was used; a maximum injection time (max IT) of 120 ms and automatic gain control (AGC) target of 10^4^ was used; ion trap CID was performed with a normalized collision energy (NCE) of 35%; the Normal scan rate setting was used; and a fragment ion matching tolerance of 0.15 Th was used. For MSX-SIM monitoring, each SIM window was 1 Th wide with an offset of +0.3 Th; each target was accumulated for up to 50 ms with an AGC target of 5×10^3^; a peak match tolerance of 10 ppm was used; the orbitrap was operated at Resolution = 60k; and an IDMS2 scan was queued if both the monoisotopic peak and the M+1 peak were matched and the monoisotopic peak exceeded a 10^4^ intensity threshold. For IDMS2 scans, CID was performed with an NCE of 35.1%; an AGC target of 10^5^ was used; for ≥ 100 targets, a max IT of 600 ms was used, whereas for < 100 targets, a max IT of 900 ms was used; the orbitrap was operated at Resolution = 15k; the fragment match tolerance was 15 ppm; and an MS3 was queued only if ≥4 fragment peaks were matched AND the cosine similarity score exceeded 0.8. Ion trap MS3 prescans were enabled. For orbitrap MS3 scans, for ≥ 100 targets, a max IT of 1,000 ms was used, whereas for < 100 targets, a max IT of 2,000 ms was used; fragment ions 50 Th below and 5 Th above the precursor m/z were excluded from SPS; 4 SPS ions were isolated for MS3 using an isolation width of 0.8 Th; HCD was performed with an NCE of 65; and the orbitrap was operated at Resolution = 50k. For experiments with dynamic close-out, close-out was triggered if a summed signal-to-noise (SumSN) of 10 * the number of channels was exceeded.

Unless otherwise specified, untargeted HRMS2 was performed with the following parameters. The expected LC peak width was 30 s. MS1 scans were performed in the orbitrap with a Standard AGC Target and max IT set to Auto, with Resolution = 60k and mass range = [400, 1600]. The Monoisotopic Peak Determination feature was used in Peptide mode. Peaks below the intensity threshold of 2.5e4 were excluded from MS2 and the Theoretical Precursor Envelope Fit feature was used with Fit Threshold = 50% and Fit Window = 0.5 Th. Only precursors with charge states in [2, 5] were included in MS2. The dynamic exclusion duration was 60 s and a mass tolerance of ±10 ppm was used for dynamic exclusion. Alternate isotopes and charge states were included in dynamic exclusion, and exclusion was enabled within the scan cycle. For MS2, a normalized AGC target of 250% was used with a max IT of 86 ms; an isolation width of 0.5 Th was used; HCD was performed with NCE = 37%; the low end of the m/z range was set to 110 Th and the high end was automatically set to 10 + precursor m/z; and the orbitrap was operated at Resolution = 50k. For untargeted HCD-HRMS2 analyses of TMTproD 35plex samples, these same parameters were used, except the orbitrap was operated at Resolution = 120k and max IT = 246 ms. For untargeted CID-HRMS2 for building the cysteine library, max IT of 150 ms was used for MS2; CID was performed with NCE = 34, CID activation time = 10 ms, and activation Q = 0.25; and the orbitrap was operated at Resolution = 15k.

Unless otherwise specified, untargeted RTS-SPS-MS3 was performed on the 2-cell-line 35plex with the following parameters. MS1 spectra used a Standard AGC target with Automatic max IT; the scan range was [350, 1500]; and the orbitrap was operated at Resolution = 120k. The Monoisotopic Peak Determination feature was used in Peptide mode. Precursors were selected for MS2 if they exceeded a 5e3 intensity threshold and if charge was in [2, 5]. Dynamic exclusion parameters were the same as HRMS2 above. For MS2, a Standard AGC target was used with max IT = 35 ms; the isolation window was 0.5 Th; CID was performed with NCE = 35% for 10 ms and activation Q = 0.25; ions were analyzed in the ion trap with an Automatic scan range and Rapid scan rate setting. Real-time search was performed on each MS2 using concatenated canonical + isoform reference proteomes from Uniprot further concatenated with contaminant proteins marked “contaminant” and further concatenated with reversed proteins marked with “##”. Cys carbamidomethylation and lysine and peptide N-terminus TMTpro labeling were included as static modifications and Met oxidation was included as a variable modification. Up to 2 missed cleavages and up to 2 variable modifications per peptide were considered. TMT SPS MS3 Mode and Close-Out were both enabled with a maximum of 2 peptides per protein and a maximum search time of 100 ms. The score thresholds were Xcorr ≥ 1.4 and dCn ≥ 0.1 with a precursor mass tolerance of 20 ppm for z = 2 and 10 ppm for z = 3 or 4. Peptides only matching proteins marked with “##” or “contaminant” were excluded from SPS-MS3. SPS ions were excluded between precursor m/z – 50 Th and precursor m/z + 5 Th and TMTpro tag loss ions were excluded from SPS. 10 SPS ions were used. For MS3, a normalized AGC target of 300% was used with a max IT of 246 ms. The peptide isolation window was 1.2 Th and the fragment isolation window was 2 Th. HCD was performed with NCE = 55% and reporter ions were analyzed in the orbitrap with scan range = [110, 1000] and Resolution = 120k.

### Data Processing and Analysis

Unless otherwise specified, GoDig data were processed in the following manner. MS3-level data were extracted from the raw files using the Data Analysis tab of the GoDig software. In R, MS3-level data were then collapsed to the precursor, peptide, or site level by summing the signal-to-noise ratios in each channel. An analyte (precursor, peptide, or site) was considered quantified only if the SumSN exceeded 10 * (# channels). For quantitative analysis, any analyte that failed to exceed this was discarded. In R, quantitative data were then corrected using the isotopic purity values contained in the certificate of analysis for the TMT reagent lot, publicly available on www.thermofisher.com.

Untargeted data were searched using a comet-based in-house pipeline. Cysteine carbamidomethylation and TMT labeling of peptide N-termini and K were set as static modifications and M oxidation was used as a variable modification; for phosphoproteomics, STY phosphorylation (79.966331 Da) was included as a variable modification with a possible neutral loss of 97.976896 Da; for ubiquitylomics, K diGly was included as a variable modification (114.0429274 Da); for SLC-ABPP, the DBIA adduct was included as a variable modification on C (239.162826 Da). The precursor m/z tolerance was 50 ppm. The fragment m/z tolerance was 1.0005 Th for ion trap MS2 and 0.02 Th for orbitrap MS2. Peptide-spectrum matches were adjusted to a 1% false discovery rate (FDR) using a linear discriminant analysis, protein inference was performed using the parsimony algorithm (Zhang, Chambers, and Tabb, *J. Proteome Res*. **2007**, *6*, 3549), and then proteins were filtered to a final protein-level FDR of 1% (Huttlin, et al., *Cell* **2010**, *143*, 1174). Sites were localized using the AScorePro algorithm (Gassaway, et al., *Nature Methods* **2022**, *19*, 1371). Quantitative data were corrected using the isotopic purity values contained in the certificate of analysis for the TMT reagent lot.

For AD vs. NCI differential analysis, a two-tailed T-test was performed assuming unequal variance, and then the *p*-values or significance threshold was adjusted using the Benjamini-Hochberg method. For dose-response analysis, a 4-parameter logistic regression was performed using the drc package in R, with the zero-dose asymptote *c* fixed to be the average of the DMSO measurements (conc = 0 µM) within that subplex (Seber, G. A. F. and Wild, C. J (1989) *Nonlinear Regression*, New York: Wiley & Sons [p. 330]). For Fig. 6D, a compound-site interaction was only counted if (a) the IC_50_ parameter *e* was < 100 µM, (b) the slope parameter *b* was positive (increasing compound dose decreased DBIA signal), and (c) the infinite-dose asymptote *d* was less than *c*/2. For visualization (Fig. 6E–F), the effect of peptide interference was removed by scaling the data such that the zero-dose asymptote was set to 1 and the infinite-dose asymptote was set to 0.

GoDig libraries were built in the GoDig software. Peptides with a chromatographic peak width (measured by in-house software) > 10 min were excluded from libraries. A fragment mass tolerance of 0.02 Th was used in library building. Indexed databases for real-time search were built in the GoDig software using the same search parameters described above and with peptide lengths in [7, 63].

### Software Development

GoDig feature implementation and debugging was done in C# using Visual Studio 2022 (Microsoft).

### Covalent Docking

Covalent protein-ligand complex structures were predicted using Schrodinger Maestro Version 13.1.141, release 2022-1.65,66,67. TXNDC12 was prepared using default settings in the protein preparation wizard with a single protein chain (chain A) and ligand. Covalent docking experiments were conducted using AlphaFold 3.0 structure AF-O95881-F1-v4. The structures underwent minimalization using default settings. E390 was prepared using default LigPrep settings and a maximum of 32 states were generated. A 20 Å box centered around residue 66 was used. Covalent docking was conducted in pose prediction mode using Cys66 of the protein as the reactive residue, Michael addition reaction SMARTS, a 3.5kcal/mol energy cutoff and a maximum output of 3 poses per ligand.

## DATA AVAILABILITY

All raw data, GDVXML files, search results, GoDig outputs, and GoDig libraries generated or used in this work have been deposited in the PRIDE (Perez-Riverol, et al., *Nucleic Acids Res* **2022**, *50*, D543) repository via the ProteomeXchange Consortium with the identifier PXD065227.

## CODE AVAILABILITY

The GoDig software is freely available via a Recipient Agreement for the Orbitrap Eclipse or Orbitrap Ascend mass spectrometer with an iAPI license. The Recipient Agreement and instructions for obtaining the iAPI license can both be obtained at https://gygi.hms.harvard.edu/software.html. GoDig is enabled through the iAPI framework such that the GoDig source code is not available, but iAPI sample code is available at https://github.com/thermofisherlsms/iapi. GoDig uses Comet (ver. 2025.02.0), available at https://uwpr.github.io/Comet.

## ACKNOWLEDGEMENTS

The authors thank members of the Gygi Laboratory for helpful conversations and support as well as the Stanford/VA/NIA Aging Clinical Research Center (ACRC) and the patients and their families for brain samples.

This work was funded in part by the National Institutes of Health (NIH) grants AG088297 (S.R.S.), GM67945 (S.P.G.), GM132129 (J.A.P.), and CA282268 (Q.Y.).

## AUTHOR CONTRIBUTIONS

S.R.S., S.P.G., and Q.Y. conceived the project. S.R.S. and Q.Y. implemented features in GoDig. S.R.S. prepared the 5-cell-line samples, the 2-cell-line sample, the human brain samples, and the cysteine library samples, performed all LC-MS/MS experiments, analyzed and plotted all data, and wrote the manuscript. G.A.F. cultured and treated cells and prepared samples for the dose-response experiments and performed *in silico* docking. C.B. and M.W.M. obtained and homogenized human brain samples. J.D.C. implemented features in iAPI. N.R.Z. provided lysates for the 2-cell-line mixture. B.M.G. and S.D. provided lysates for the 5-cell-line mixture. K.H.O. aided in code optimization. J.A.P. assisted with instrument maintenance. S.R.S., G.A.F., J.A.P., S.P.G., and Q.Y. edited the manuscript.

