## Supporting Information for "Next-Generation Multiplexed Targeted Proteomics Quantifies Post-Translational Modifications, Compound-Protein Interactions, and Disease Biomarkers with High Throughput"

### Figures S1–S9

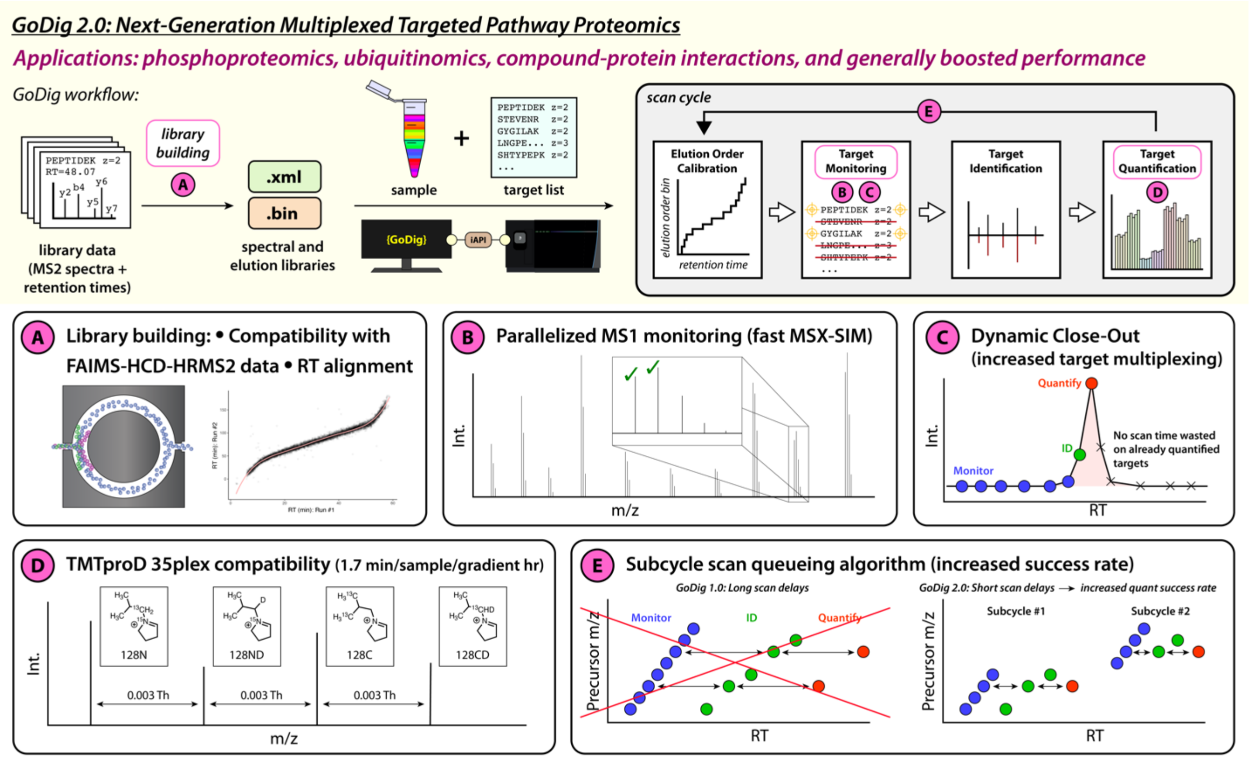

**Figure S1.** Detailed overview of the improvements made in GoDig 2.0.

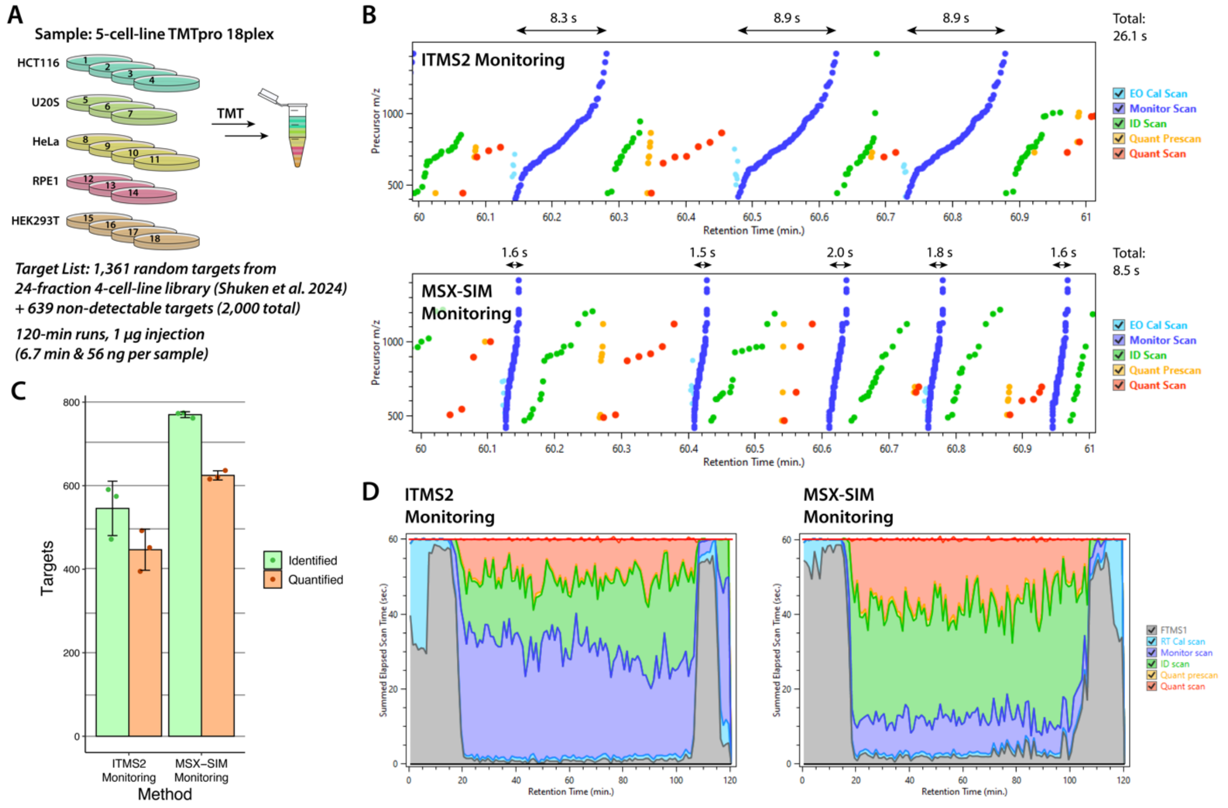

**Figure S2. MSX-SIM monitoring increases the number of quantifiable targets by monitoring more quickly. A.** Sample and target list used for GoDig 2.0 feature testing. **B.** GoDigViewer (Ref. 9) plot of typical runs targeting the 2,000 randomly selected targets with ion trap MS2 (ITMS2) and MSX-SIM monitoring. The x-axis is zoomed in on scans between RT = 60 min and 61 min. EO Cal = elution order calibration. Quant Prescan = short MS3 performed in ion trap for calculating injection time for Quant Scan (longer MS3 performed in orbitrap). **C.** Total targets identified and quantified in runs performed in triplicate. **D.** Stacked area plots from GoDigViewer representing amounts of time spent on each type of scan during a typical run with each method. The Elapsed Scan Time was calculated for each scan by subtracting the RT of the scan from the RT of the following scan. Summed Elapsed Scan Time is the sum of these times during one minute of acquisition. Orbitrap MS1 (FTMS1) scans are conducted between scan cycles.

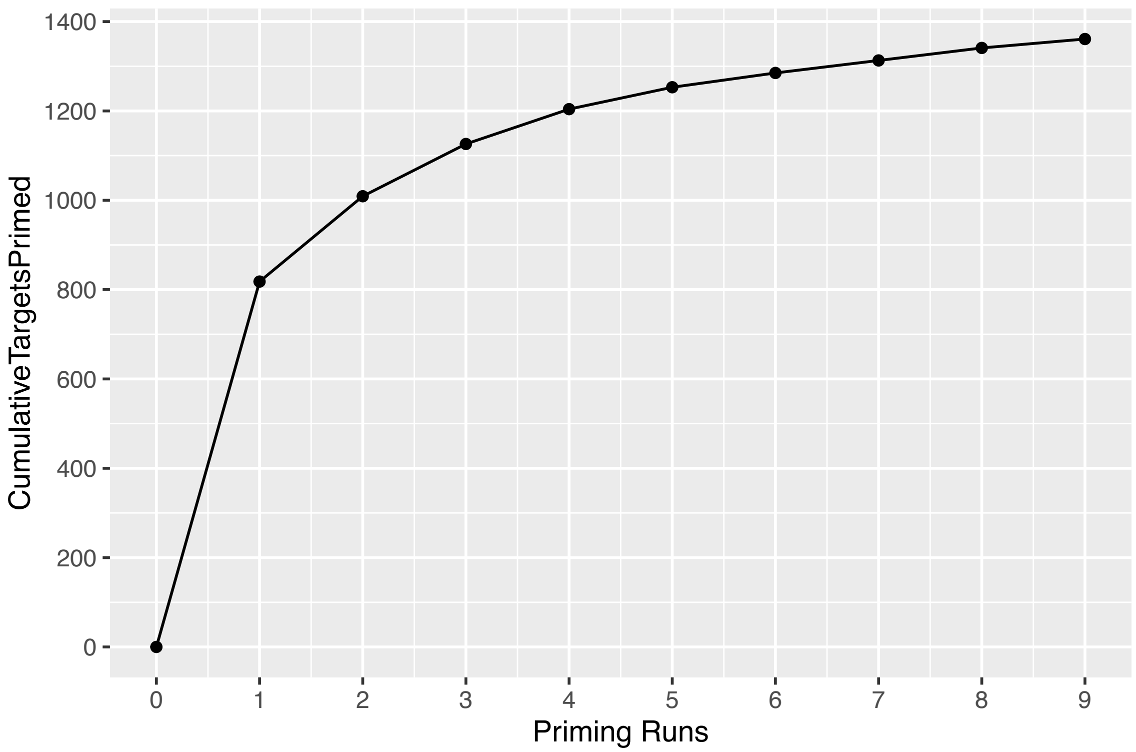

**Figure S3.** Targets from a list of 2,000 random targets primed (identified by IDMS2) in the 5-cell-line mixture (Fig. S2A) during 9 priming runs performed as described in Shuken et al. 2024.

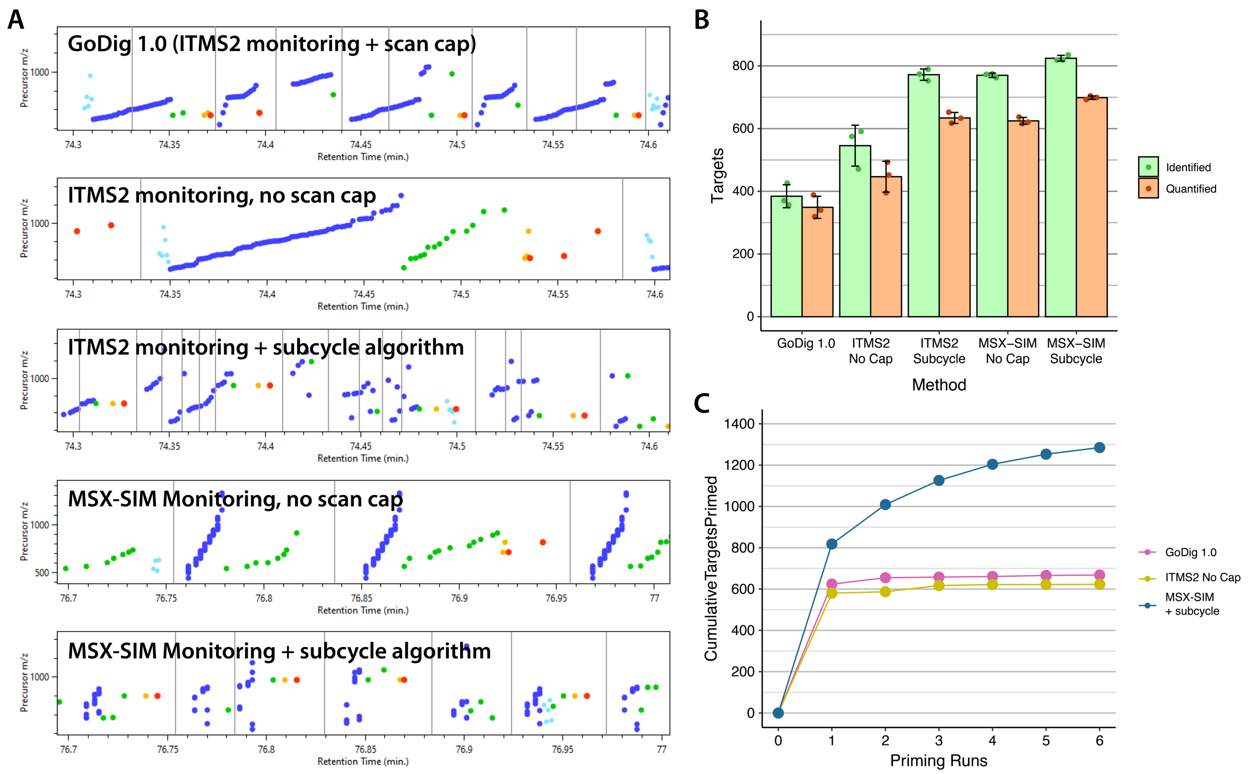

**Figure S4. The subcycle scan queueing algorithm increases performance by reducing scan delay. A.** GoDigViewer visualizations of typical runs performed with each of the five indicated methods, zoomed in to an RT range 0.3 min wide. Color code is in Fig. S2B. **B.** Identified and quantified targets in triplicate runs with these five methods targeting 2,000 randomly selected targets. **C.** Results of 6 priming runs performed with GoDig 1.0 (ITMS2 monitoring + scan cap), ITMS2 monitoring without scan cap, and MSX-SIM with the subcycle algorithm.

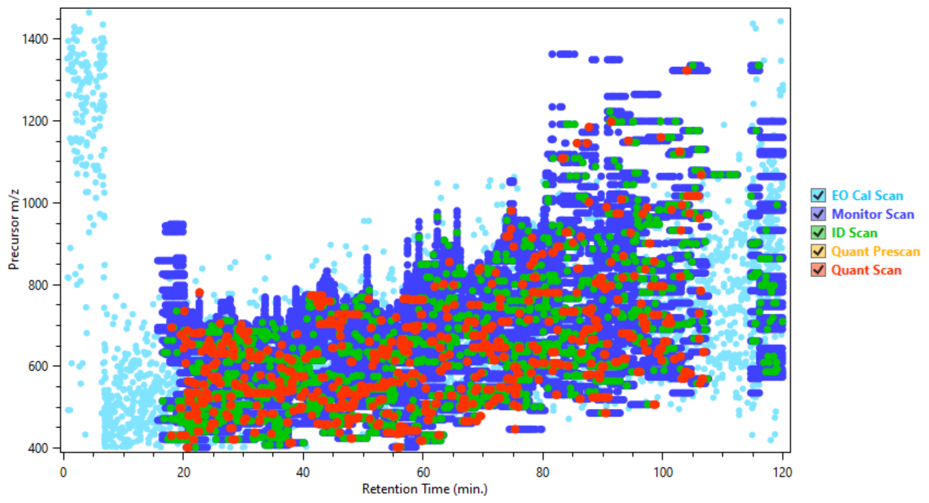

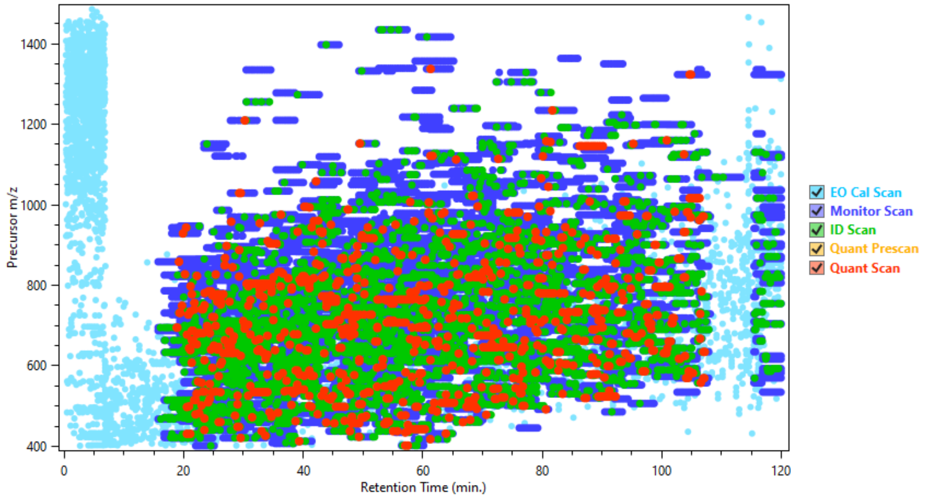

**Figure S5. GoDigViewer visualization of 2,000 random precursors targeted in 5-cell-line mixture with GoDig 1.0 and with the monitor scan cap removed.** Top: GoDig 1.0. Bottom: GoDig 1.0 with scan cap removed (the same experiment depicted in Fig. S2B, top).

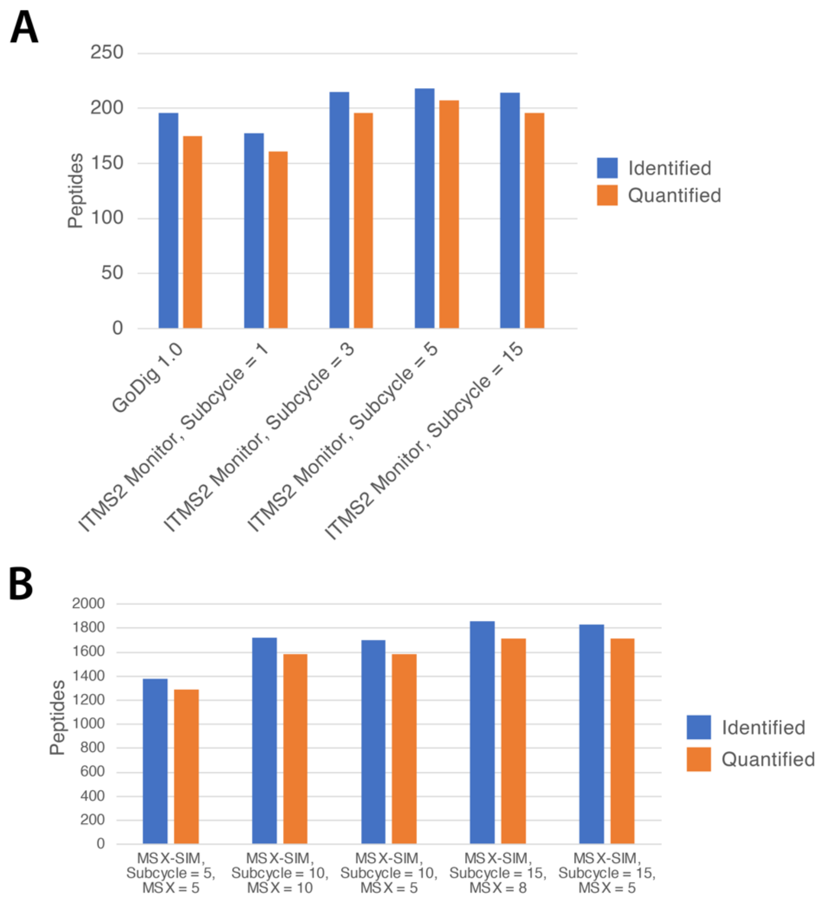

**Figure S6. Performance comparisons of subcycles with different subset sizes. A.** Runs targeted 400 random targets. **B.** Runs targeted 2000 abundant peptides. "Subcycle" = subset size as described in the main text. "MSX" = number of targets per MSX-SIM scan.

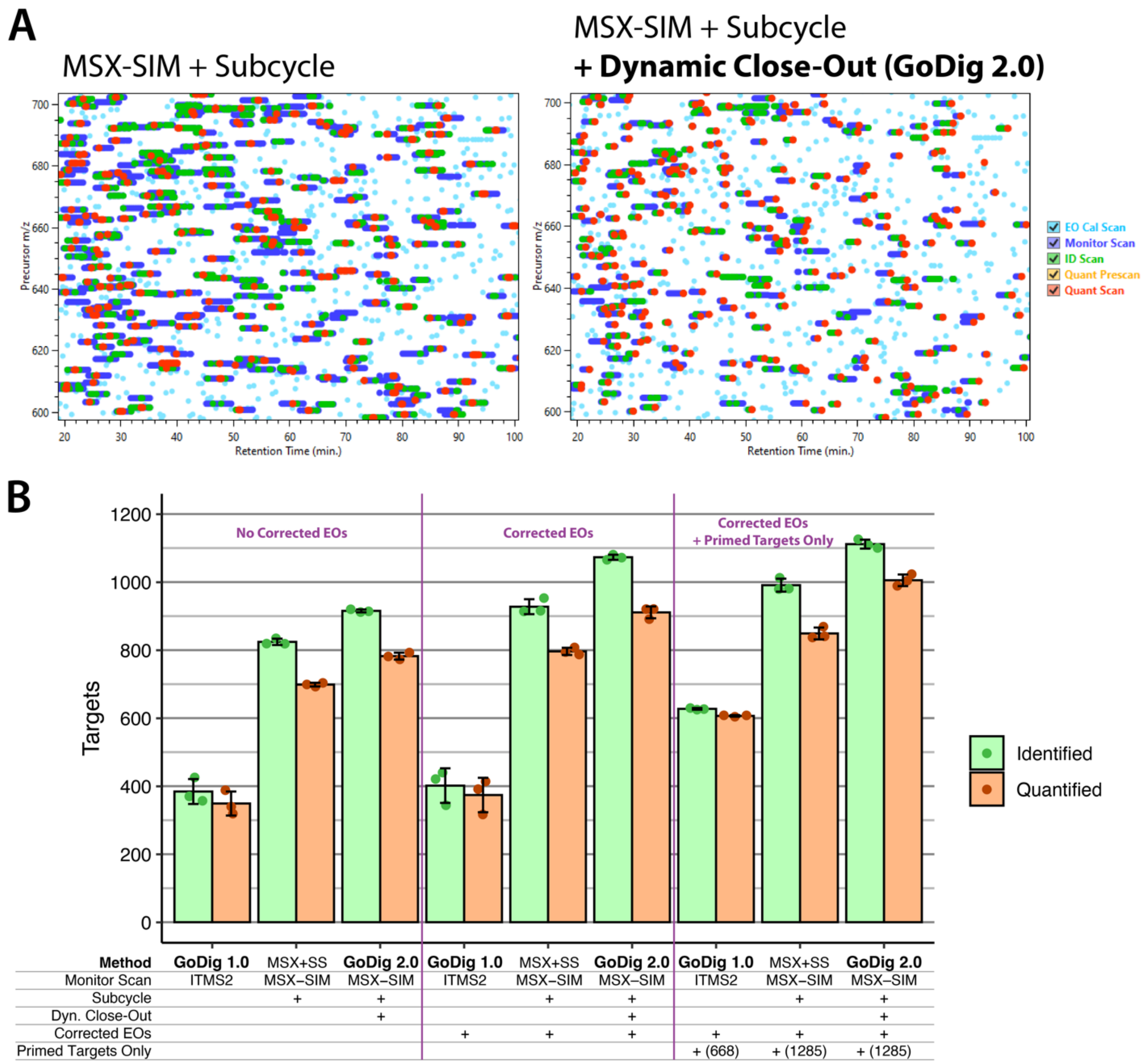

**Figure S7. Dynamic close-out increases success rates by liberating scan time. A.** GoDigViewer screenshots of typical runs using MSX-SIM monitoring and the subcycle algorithm with and without dynamic close-out enabled. **B.** Comparisons between GoDig 1.0, GoDig 2.0, and GoDig with MSX-SIM monitoring and the subcycle algorithm without dynamic close-out ("MSX+SS"). Corrected EOs = using elution order (EO) information from priming runs. Primed Targets Only = restricting target list to targets chromatographically primed during priming runs.

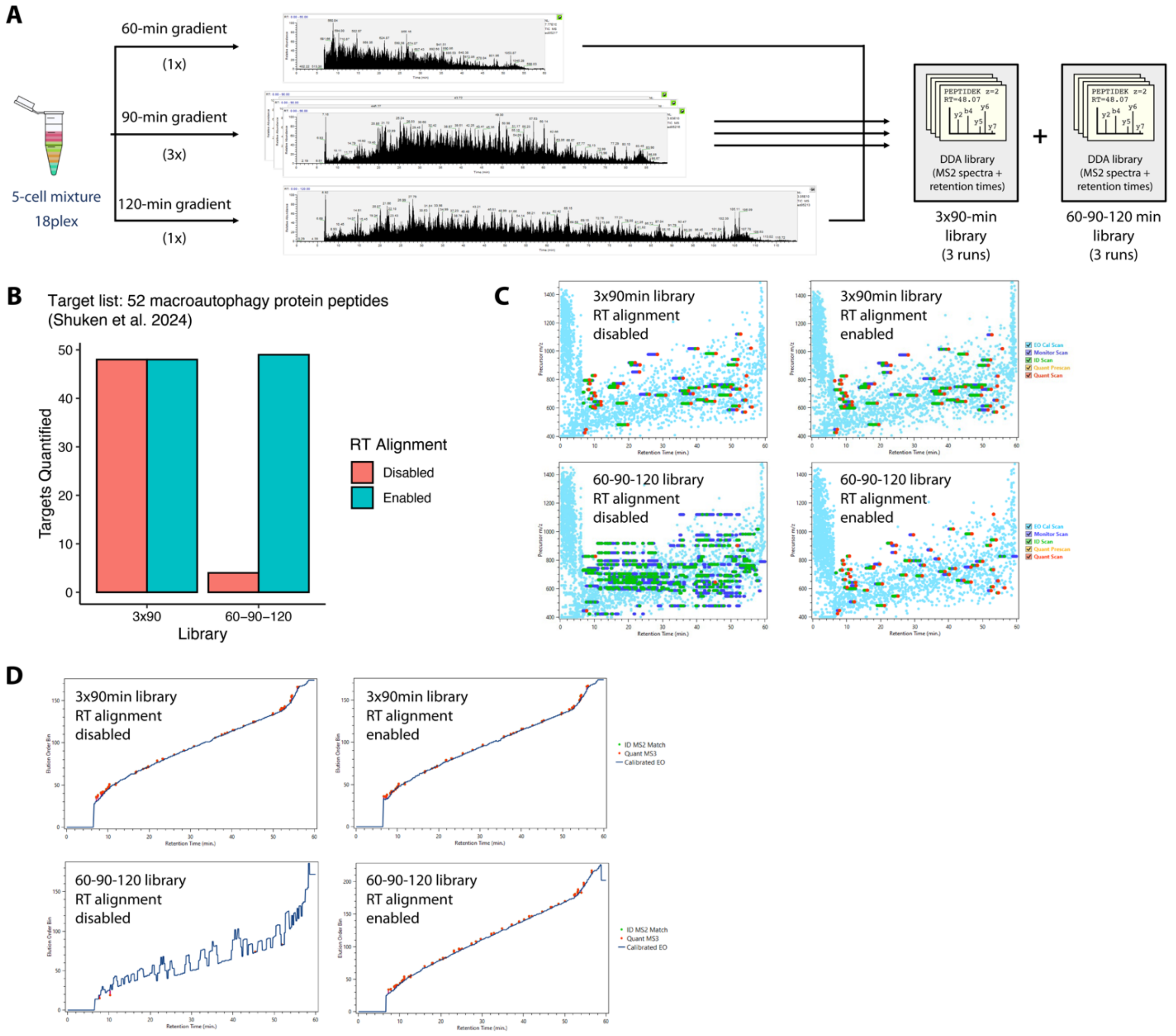

**Figure S8. Retention time (RT) alignment in library building. A.** The RT alignment feature was tested by building libraries from 3 runs with or without different gradients (and gradient lengths), with or without the feature enabled. **B.** RT alignment recovers performance when targeting 52 peptides from macroautophagy proteins present in these libraries (Shuken et al. 2024). **C.** GoDigViewer visualization of GoDig runs under these 4 conditions. **D.** Visualization of EO calibration under these 4 conditions. ID MS2 Match = successful IDMS2. Calibrated EO = current calibrated EO bin.

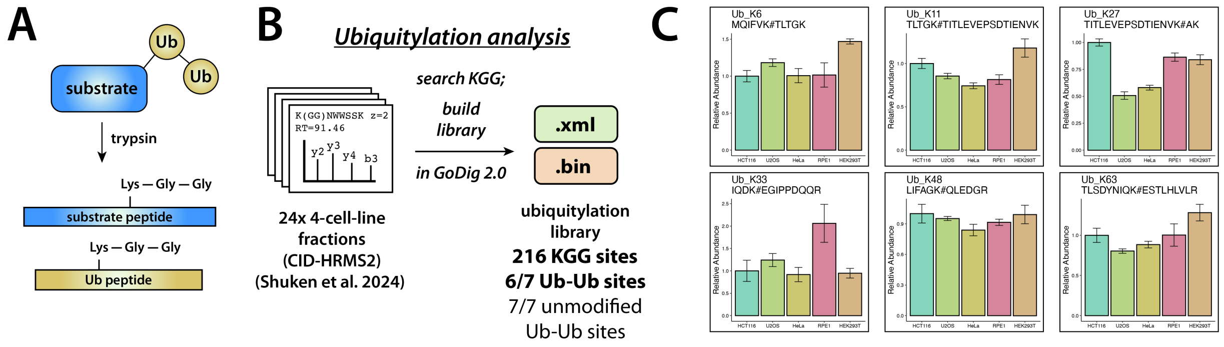

**Figure S9. Polyubiquitin branching analysis with GoDig 2.0. A.** Diagram of Lys-Gly-Gly (KGG) modification resulting from tryptic digestion of proteins post-translationally modified with ubiquitin (Ub) and polyubiquitin (polyUb). **B.** Construction of spectral and elution libraries containing 216 ubiquitylation sites (Lys-diGly, KGG) from published 4-cell-line fraction data in GoDig 2.0. In addition to 6/7 Ub-Ub branching sites, the library contains unmodified peptides that cover all 7 unmodified lysines. C. Quantification of 6 (out of 7 known) ubiquitin-ubiquitin branch sites with GoDig 2.0.

### Explanations of Technical Improvements to the GoDig Platform

#### Sequential Accumulation-MS1 (MSX-SIM) Monitoring Increases Success Rates and Target Throughput by Increasing Monitor Scan Speed

One source of failure to quantify targets using GoDig 1.0 is time efficiency; to address this, we developed sequential accumulation-MS1 (MSX-SIM) monitoring (Fig. S1B). In MSX-SIM scans, each target precursor is isolated by the quadrupole (using an isolation width of, e.g., 1.0 Th) and accumulated, and then the precursors are simultaneously analyzed in the Orbitrap. “MSX” refers to the sequential accumulation and then simultaneous analysis of different scan ranges and “SIM” (single-ion monitoring) refers to the narrow isolation windows and MS1-level analysis.

When GoDig establishes which targets are within range of the current elution order (EO) bin, monitor scans for the in-range targets are queued. In GoDig 1.0, monitor scans are ion trap CID-MS2 scans; if 6 peaks match the m/z values of fragments predicted for the target, an identification (ID) MS2 scan is queued for that target. Because a typical injection time for a Monitor MS2 scan is 50 ms and ion trap CID fragmentation takes 10 ms, monitoring ten targets with GoDig 1.0 will typically take about 600 ms; by contrast, because the injection times associated with SIM scans are shorter and no fragmentation is performed, most MSX-SIM scans take about 128 ms—the duration of the orbitrap transient at 60k resolution—even for up to ten targets.

Fig. S2B shows an illustrative example of how MSX-SIM monitoring improves speed relative to ITMS2 monitoring: in this time period, monitoring in-range targets took >8 s with MS2 and <2 s with MSX-SIM, enabling more scan cycles to occur, and therefore more IDMS2 and MS3 scans. As shown in Fig. S2D, this results in a ~3x reduction in total time spent monitoring throughout runs. This saved time is spent identifying and quantifying targets; when performed in triplicate, MSX-SIM monitoring yields 39.9% more successful quantifications (Fig. S2C).

#### The Subcycle Algorithm Reduces Scan Delay and Enables Priming of 94.4% of Randomly Selected Challenging Targets

Another source of failure to quantify targets with GoDig 1.0 is scan delay, i.e., the time between a given target’s successful monitor scan and the subsequent IDMS2, or between IDMS2 and MS3 (Fig. S1E). To reduce scan delay, GoDig 1.0 inherently limits the number of targets that can be monitored at once—i.e., a hard cap of 15 targets per cycle—which helps reduce scan delay, but limits how many targets can be quantified in a run, consistently reducing performance

(Fig. S4A–C). (The experiment described in the previous section and depicted in Fig. S2 was performed with the scan cap removed; Fig. S5 illustrates the same experiment with the scan cap applied.) We implemented a new alternative that targets all precursors while reducing scan delay, which we call the subcycle algorithm (Fig. S1E). In this scheme, the monitor scans for a subset of the in-range targets are submitted, and the subsequent ID MS2s and MS3s are executed, all before the next subset is submitted. Subsets should be kept as small as possible without sacrificing performance: for ITMS2 monitoring, a subset size of 5 targets is optimal, whereas for MSX-SIM monitoring, the optimal subcycle size is 15 targets, with 5 targets monitored per MSX-SIM scan (Fig. S6).

Removing the monitor scan cap from GoDig 1.0 enables the quantification of targets that would be excluded by the cap and increases success rates, but the increase in scan delay still limits successful quantifications (Fig. S4A–B). Both with ion trap MS2 (ITMS2) monitoring and MSX-SIM monitoring, the subcycle algorithm significantly increases the quantification success rate, giving a 12% boost with MSX-SIM, a 42% improvement over ITMS2 without the scan cap, and an 82% boost over GoDig 1.0 (Fig. S4B). This is exemplified in Fig. S4A, in which the scan delay is reduced for several targets. The subcycle algorithm also enabled the chromatographic priming of 1,285 (94.4%) of the 1,361 detectable targets over 6 runs (Fig. S4C), a >2x improvement over GoDig 1.0, which becomes overwhelmed as soon as the EO tolerance widens to ±15 bins (±7.5 min), resulting in the priming of only 623 (45.8%).

#### Dynamic Close-Out Increases Success Rates by Improving Time Efficiency

To further improve time efficiency, we implemented a feature we call dynamic close-out, with which GoDig stops monitoring a target once it has been quantified (i.e., the cumulative summed TMT reporter ion signal-to-noise ratio has exceeded 10 per channel) (Fig. S1C). By avoiding performing monitor and ID scans on peptides that have already eluted, more time is available for analyzing more targets (Fig. S7A).

We call the simultaneous employment of MSX-SIM monitoring, the subcycle scan queueing algorithm, and dynamic close-out "GoDig 2.0." With all of these features enabled, GoDig 2.0 was able to quantify 782 ± 10 precursors in a single run, compared to only 349 ± 35 for GoDig 1.0—a 2.2x improvement (Fig. S7B). Of the 433 quantification successes, 84 (19%) of these came from dynamic close-out. Using the corrected EO bins determined by the priming runs illustrated in Fig. S4C, GoDig 2.0 quantified 911 ± 17 targets (115 due to dynamic close-out), 2.4x more targets than GoDig 1.0.

The deeper priming made possible by MSX-SIM monitoring and the subcycle algorithm allowed the establishment of a new target list: the 1,285 primed targets. Using the corrected EO bins from priming as well as this trimmed target list, GoDig 2.0 was able to quantify 1005 ± 17 targets in a single run (156 from dynamic close-out) compared to only 606 ± 2 with GoDig 1.0 using the 668 targets primed with GoDig 1.0 (Fig. S7B).
